# Low-barrier hydrogen bond determines target-binding affinity and specificity of the antitubercular drug bedaquiline

**DOI:** 10.1101/2023.07.28.551034

**Authors:** Joanna Słabońska, Subrahmanyam Sappati, Antoni Marciniak, Jacek Czub

## Abstract

The role of short strong hydrogen bonds (SSHB) in ligand-target binding remains largely unexplored, thereby hindering a potentially important avenue in the rational drug design. Here, we investigate the interaction between bedaquiline (Bq), a potent anti-tuberculosis drug, and the mycobacterial ATP synthase, to unravel the role of a specific hydrogen bond to a conserved acidic residue in the target affinity and specificity. Our ab initio molecular dynamics simulations reveal that this bond belongs to the SSHB category and accounts for a substantial fraction of the target binding energy. We also demonstrate that the presence of an extra acidic residue (D32), found exclusively in mycobacteria, cooperatively enhances the HB strength ensuring the specificity for the mycobacterial target. Consistently, we show that the removal of D32 markedly weakens the affinity, leading to Bq resistance associated with mutations of D32 to non-acidic residues. By designing simple Bq analogs, we then explore the possibility to overcome the resistance and potentially broaden the Bq antimicrobial spectrum by making the SSHB independent on the presence of the extra acidic residue.

---

Hydrogen bonds (HB) are ubiquitous intermolecular interactions characterized by directionality and specificity. Owing to their versatility and robustness, HBs are essential for many physical phenomena, from protein structure and function to molecular recognition and self-assembly to chemical reactivity.^1,2^ As hydrogen-bond energies typically vary in the 2–10 kcal/mol range, HBs are generally considered weak non-covalent interactions.^3,4^ For these weak HBs, the distance between the donor (D) and acceptor (A), d_DA_, is typically *>*2.6 Å and the proton (H) is largely localized on the donor,^5,6^ which corresponds to a strongly asymmetric double-well potential with a more pronounced minimum at a short D–H distance (purple in Fig. 1A).^7^

**Figure 1.**
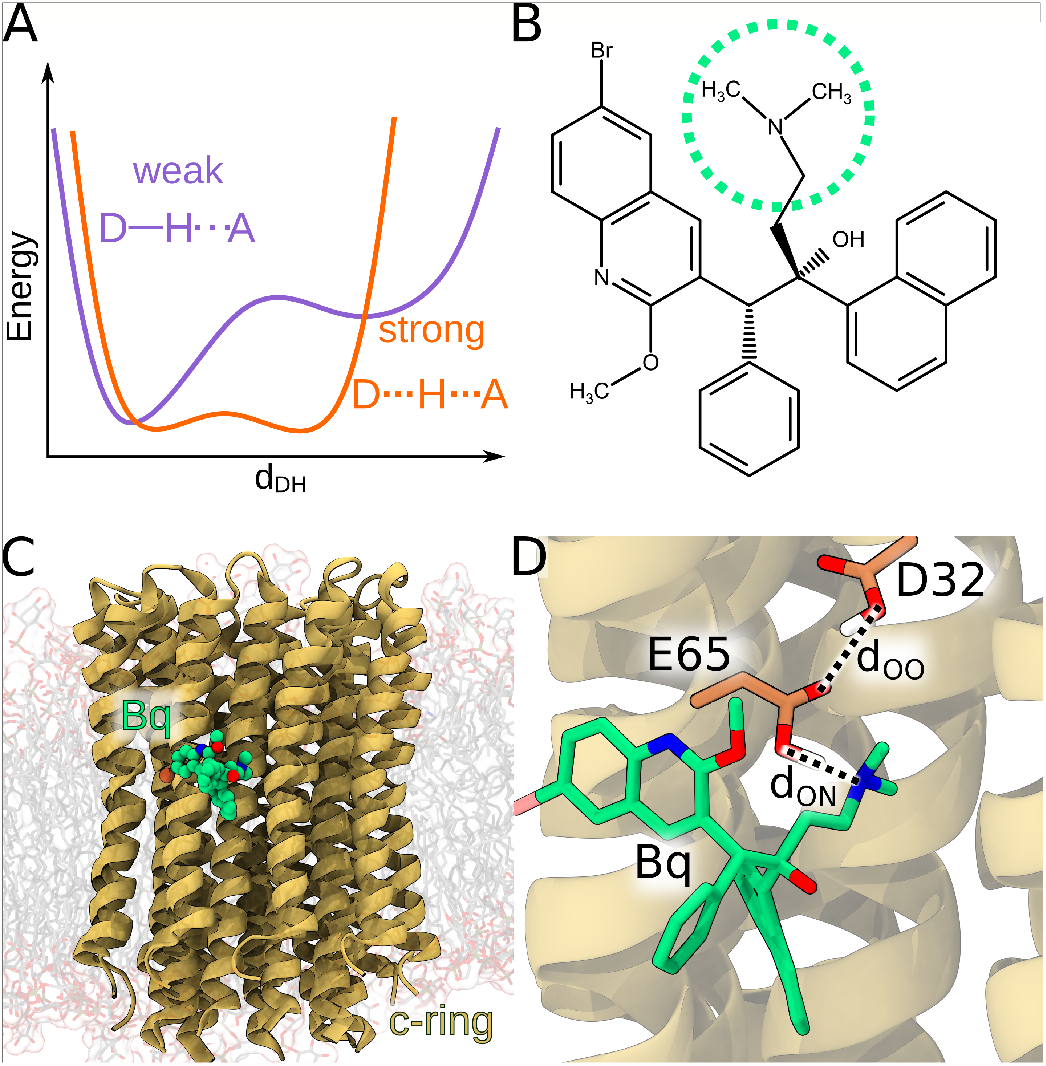
(A) Schematic view of the energy landscape for the displacement of the H-bonded proton between the donor, D, and the acceptor, A, for the weak (purple) and strong (orange) HB. (B) Chemical structure of bedaquiline (Bq). Tertiary amino group forming involved in the hydrogen bonding interaction with the c-ring (yellow) is indicated by the green circle. (C) Side view of the c-ring with bound Bq. (D) Hydrogen bond between the Bq amino group and the c-ring E65. Unique to the mycobacterial c-ring is the presence of an additional acidic residue, D32, which, due to its location, can form an additional HB with E65. For more structural details of the Bq–c-ring interface, see Fig. S1.

However, if D and A have similar proton affinities (or pKa’s), HB energy may greatly exceed the normal range occasionally reaching 25–40 kcal/mol, while d_DA_ becomes markedly shorter (2.5 Å or even less).^7–10^ In such cases, the proton experiences a more symmetric low-barrier or single-minimum potential (orange in Fig. 1A) and the HB exhibits some covalent character, as evidenced, e.g., by topological analysis of electron density.^6,11^ If such short strong HBs (SSHB) form in nonpolar environments, the absence of competing hydrogen bonding to solvent make their free energy of formation particularly favorable.^6,12,13^

So far, the role of unusually strong HBs in ligand–target binding has been largely unexplored, and thus little attention has been paid to such interactions in medicinal chemistry.^14,15^ To show that SSHB can in fact be utilized in structure-based drug design, here, we demonstrate that a specific strong HB is critical for the target affinity and specificity of the important anti-tubercular drug bedaquiline (Bq) and that weakening of this bond due to a mutation is responsible for drug resistance. We also point out potential ways to combat this resistance mechanism and/or broaden the spectrum of activity of Bq to other bacterial pathogens.

Bq (Fig. 1B) is a potent diarylquinoline inhibitor of the mycobacterial ATP synthase that has been used for almost 10 years against multi- and extensively drug-resistant tuberculosis.^16,17^ X-ray and cryo-EM structures have revealed that Bq binds to the outer surface of the c-ring – the rotary part of the membrane-embedded F_o_ subcomplex of ATP synthase (Fig. 1C).^18,19^ Despite some degree of shape complementarity, (Fig. S1B), mostly non-polar interactions between Bq and the shallow binding pocket on the c-ring (Fig. S1C) cannot alone account for the experimental binding free energy of ∗ 8 kcal/mol,^20,21^ especially in the lipid environment where the ligand–protein complex cannot be stabilized by hydrophobic attraction. Apparently, the remaining driver is the only polar contact, i.e., the hydrogen bond between the amino group of Bq (encircled in Fig. 1B) and the conserved c-ring glutamate (E65 in Fig. 1D). Indeed, the Bq/c-ring crystal structure indicates that this bond is unusually short (with the average d_DA_ = 2.4 Å),^18^ and hence it likely falls within the SSHB category. The fact that binding mechanism of Bq to the c-ring is not fully understood hinders the development of its novel derivatives that would evade the emerging resistance^22–24^ or target ATP synthases other than the mycobacterial one.^25,26^

To elucidate this mechanism, and, particularly, the role of the short HB in the target affinity and selectivity of Bq, we initially determined the free energy landscape governing the Bq binding to the membrane-embedded c-ring, using force field-based molecular dynamics (MD) simulation (green in Fig. S2).

The binding free energy (Δ*G*) obtained in this way (ca. −1 kcal/mol) turned out to be much smaller than that measured by surface plasmon resonance (−8 kcal/mol),^20^ even though the simulated structure of the complex closely matches the experimental one (RMSD of 0.22 nm; see Fig. S3). In fact, the computed affinity was similarly low even when the HB was prevented by removing the proton from E65 (red in Fig. S2). Thus, we concluded that the classical description is not adequate to capture the contribution due to the short HB and that the remaining Bq–c-ring interactions contribute relatively little to the target affinity.

Therefore, to understand the nature of the short HB stabilizing the Bq/c-ring complex, we turned to a hybrid QM/MM ab initio MD approach based on density functional theory as implemented in the NAMD/Orca interface^27^ (for the definition of the QM regions and other details, see Fig. S4 and SI Methods). Specifically, we determined the effective HB potential similar to those in Fig. 1A, by computing how the free energy changes with the oxygen–proton distance, d_OH_, using the umbrella sampling method (see SI Methods). The resulting profile (Fig. 2B) shows two free energy wells corresponding to the proton residing on the carboxylate O atom of E65 or the amino N atom of Bq. As can be expected for the low dielectric of the lipid bilayer, the “salt-bridge” configuration with the proton on Bq is slightly disfavored (by ∼0.8 kcal/mol) compared to the neutral configuration with the proton on E65. Importantly, the barrier for the proton transfer between the two heteroatoms (1 kcal/mol) is markedly lower than the typical zero point energy in the O–H or N–H stretching modes (5–6 kcal/mol), implying a large degree of proton sharing between O and N which is indicative of the formation of SSHB. Consistently with this conclusion, the O–N distance, d_ON_, is on average 2.54 Å (Fig. 2C) and the MD ensemble-averaged energy of the HB in the neutral configuration, approximated at the DFT level and corrected for basis set superposition error, is in the range− (19−21) kcal/mol (Table S1, Fig. S8). As expected, the bond becomes particularly short (2.48 Å) and strong (−32 kcal/mol) when the proton is roughly equidistant from both heteroatoms (Fig. 2C, Fig. S7 and Fig. S8). Also, the average value of electron density at the HB critical point of 0.10 e/A^3^ is characteristic of quasi-covalent hydrogen bonds, which are usually in the range of 0.08–0.14 e/A^3^ (Fig. S9).^28^

**Figure 2.**
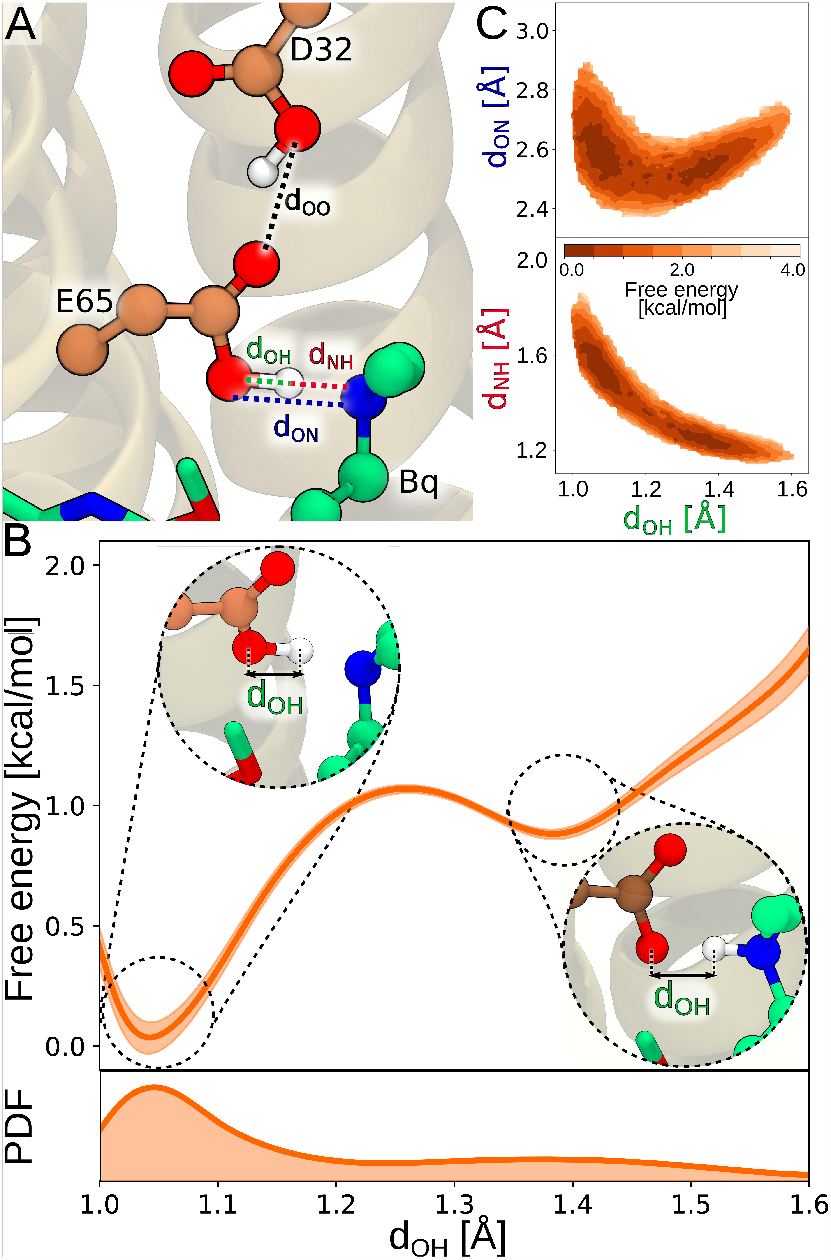
(A) Definition of the interatomic distances considered in this work. (B) Free energy profile for the proton transfer between the heteroatoms involved in the Bq–E65 hydrogen bond computed using US and the double-zeta basis set. The lower panel shows the corresponding probability distribution of the d_OH_ distance. For comparison with the profile obtained using the unbiased MD and triple-zeta basis set, see Fig. S5. Fig. S6 shows the convergence of the free energy profile. (C) Free energy landscapes describing the correlation between *d*_OH_, *d*_ON_ and *d*_NH_. For other correlations see Fig. S7.

The same conclusion may be drawn from the NBO-based analysis of the overlap between the lone-pair orbital of the acceptor, n_*A*_, and the antibonding orbital of the D–H bond, 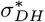 orbital. “Interaction” energy between these orbitals as a function of the proton position (Fig. S10) reach the value of 130 kcal/mol for the proton equidistant from the heteroatoms, which falls in the range characteristic for the strong HBs.^1^

In essence, our data strongly indicate that Bq forms a short low-barrier hydrogen bond with E65 on the mycobacterial c-ring, which accounts for a significant portion of its affinity for the target. Notably, our QM/MM simulations also revealed that while bound to Bq, E65 also forms a stable H-bond with an additional acidic residue, aspartate D32, which is conserved across mycobacteria (Fig. 1D). As indicated by the critical point analysis (Fig. S9), this additional interaction itself is a relatively weak HB (d_OO_ =2.65 Å), but its presence may be crucial for strengthening the Bq–E65 bond. This is because it is known that carboxyl groups are particularly prone to form SSHB when they participate in cooperative hydrogen bonding networks.^29^ To put this hypothesis to the test, we removed the additional carboxylate by replacing D32 with alanine and then used QM/MM umbrella sampling to recompute the free energy profile characterizing the Bq–E65 bond. Fig. 3A clearly shows that the profile undergoes a striking change upon D32A mutation, assuming the shape that is characteristic of a normal-strength HB, with a pronounced proton localization on the donor, which is the carboxylate of E65. In line with these findings, the average energy of the bond is reduced by ∼6 kcal/mol (Fig. S8), while the electron density at the bond critical point decreases to 0.06 e/A^3^, a value typical for a normal-strength HB (Fig. 3C and Fig. S9).

**Figure 3.**
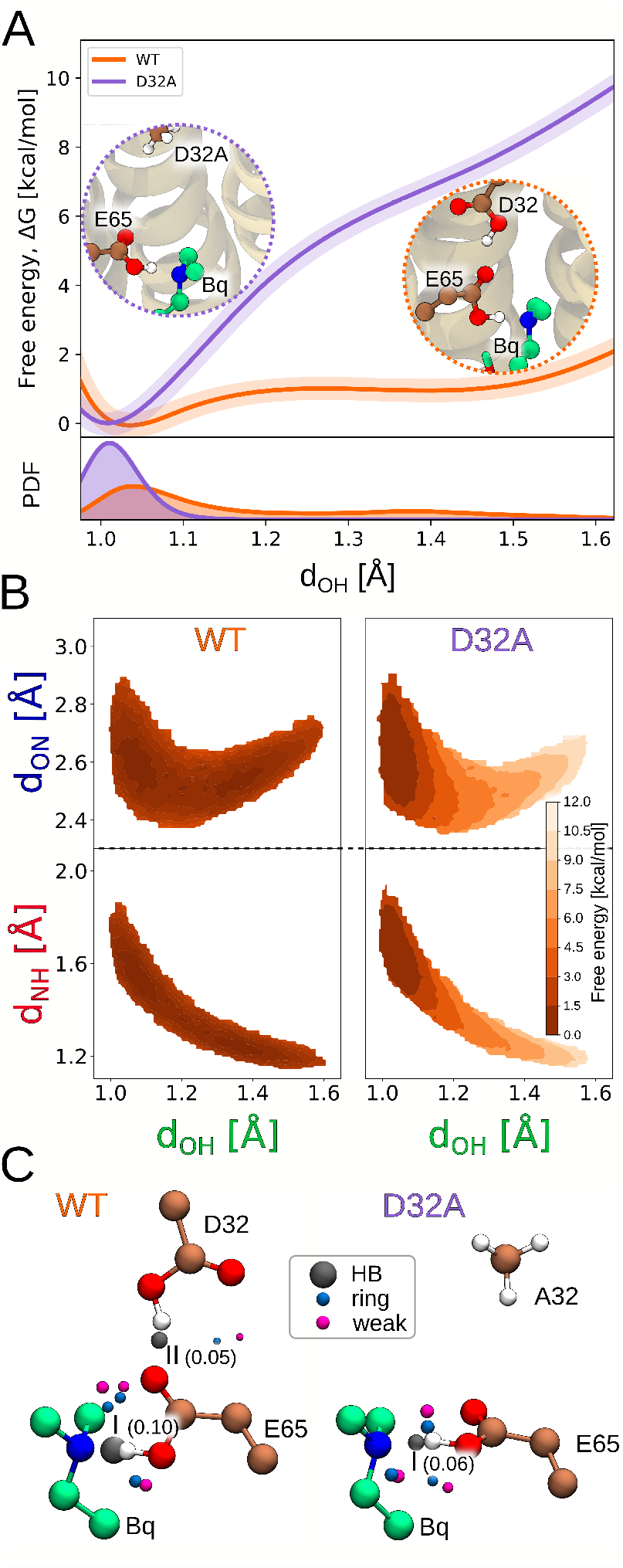
(A) Comparison of the free energy profiles characterizing the Bq–E65 HB and the corresponding probability distributions between the wild-type c-ring (WT) and its D32A variant. For convergence of the free energy profiles, see Fig. S6 and Fig. S11. (B) Free energy landscapes describing the correlation between *d*_OH_, *d*_ON_ and *d*_NH_ for Bq bound to WT and D32A. For other geometric correlations, see Fig. S12. (C) Critical point analysis characterizing the strength of interaction between the Bq amino group and E65 in the WT c-ring (left) and its D32A variant (right). For numeric data see Fig. S9.

At the same time, the average d_ON_ distance increases from 2.54 to 2.67 Å (Fig. 3B) and the HB becomes slightly less co-linear and orientationally restrained (Fig. S12). The above comparison of WT and the D32A mutant (Fig. 3) clearly demonstrates that the extra HB to the aspartate is critical for the formation of the strong low-barrier HB between Bq and E65, which in turn appears to be indispensable for high-affinity binding of Bq to the c-ring. More broadly, this is a unique example of a strong cooperativity between two H-bonded carboxylic groups that increases the strength of an intermolecular HB by more than 7.0 kcal/mol.^29^ Given that the additional conserved aspartate residue (here, D32) occurs exclusively in the mycobacterial c-ring, it can be further proposed that it is necessary for the Bq target specificity and hence anti-tubercular activity. Consistently with this line of reasoning, recent studies have associated the resistance to Bq in clinical isolates of *Mycobacterium tuberculosis* and *Mycobacterium smegmatis* with a mutation of the aspartate residue in the c-ring to non-acidic amino acids (valine, glycine and alanine).^22,30,31^ Previous attempts to enhance the therapeutic potential of Bq and widen its antibacterial spectrum have mostly focused on modifications to the diarylquinoline scaffold, particularly the naphthyl and phenyl moieties, but with only moderate success.^24,26,32,33^ However, in light of our current results, much more attention should be paid to the hydrogen bond with the conserved acidic residue (here, E65), which provides most of the affinity. Indeed, ensuring strong character of this bond in a way that is independent of the presence of the additional carboxylate (here, D32) should allow to evade the emerging resistance and, potentially, extend the application of diarylquinoline agents to other important Gram(+) and Gram(-) pathogens.

As a proof-of-concept of this approach, we designed two analogs of Bq with the amino group replaced by either 2-hydroxy-1-methyl-imidazole (I) or formamidic acid (II) substituents, that were intended to provide stronger H-bonding interaction with E65. As can be seen in Fig. 4, the binding energy of compound I is indeed considerably larger than the one computed previously in the same way for Bq (by 9.7 and 8.0 kcal/mol for WT and D32A variant of the c-ring, respectively). This enhanced affinity, even in the absence of the additional carboxyl group, can be attributed to intramolecular cooperativity where the stronger bond to E65 formed by the imidazole N3 atom (average *d*_*ON*_ = 2.48 Å) is further strengthen by the weaker bond between the 2-OH group and the second O atom of E65 (*d*_*OO*_ = 2.76 Å; see Fig. S13). Indeed, when the cooperativity is abolished by removing the 2-OH group, the bond formed by N3 is weakened (*d*_*ON*_ = 2.58 Å) and, consequently, the binding energy is reduced by up to 50 % (compound III in Fig. S14). In contrast, the compound II, although it can also form two H-bonds to E65, is characterized by markedly lower binding energy than the original drug (however, largely independent on the presence of D32). The observed decrease in affinity seem to result from enhanced conformational flexibility of the imide substituent, leading to reduced cooperativity. This emphasizes the importance of fine-tuning of the ligand H-bonding properties.

**Figure 4.**
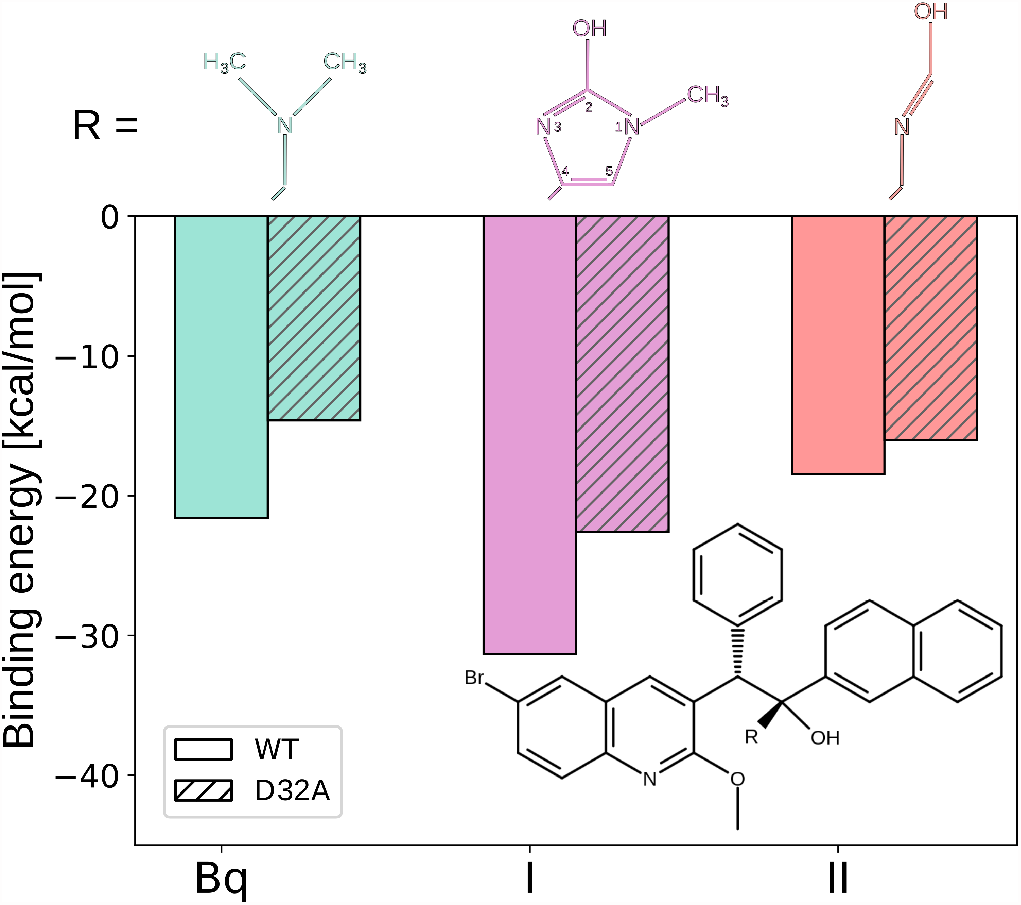
Hydrogen bond strength between Bq or its two analogs (I and II) and E65 in the WT c-ring and its D32A variant, quantified as the binding energy between the ligand’s HB-forming group (R) and either E65/D32 pair (WT) or E65 alone (D32A).

To sum up, in this work, we used ab initio and force-field-based MD simulation to show that bedaquiline (Bq), an important anti-tubercular drug, forms a short strong (low-barrier) hydrogen bond with the conserved carboxylate residue in the c-ring of mycobacterial ATP synthase (E65 in *M. phlei*), and that this bond accounts for a significant portion of Bq affinity for the target. We further demonstrated that the unique additional carboxylate residue in the mycobacterial c-ring (D32) cooperatively strengthens this HB, providing a mechanistic explanation of the target specificity. Indeed, when this additional carboxylate is removed, the Bq–E65 is weakened by ∼32 % which possibly is responsible for the emergence of the Bq-resistant isolates with D32 mutated to non-acidic residues. Building upon these findings, we proposed a novel strategy for overcoming the resistance of mycobacteria to Bq and potentially for broadening its spectrum to other bacterial pathogens. Specifically, we show that the simple imidazole analog of Bq, that, regardless of the presence of D32, preserves the strong nature of the HB to E65, has markedly higher affinity for both the WT c-ring and its D32A variant. More broadly, our results underscore the importance of considering short strong HBs in drug design and resistance mechanisms, pointing to exciting prospects in the development of novel therapeutics.

## Acknowledgement

This work was funded by the Polish National Science Centre under Sonata Bis grant No. 2017/26/E/NZ2/00472. SS wishes to acknowledge the funding provided by the Gdansk University of Technology through the Nobelium grant (No. 16/2021/IDUB/I.1) under the ’The Excellence Initiative - Research University’ (IDUB) program. This research was supported in part by PL-Grid Infrastructure. Computational resources were also provided by the TASK, WCSS and ICM Centers.

## Supporting Information Available

Detailed description of computational methods additional analyses of unbiased and free energy simulations; results obtained with different functionals and basis sets

## References

(1) Grabowski, S. J. What Is the Covalency of Hydrogen Bonding? Chemical Reviews 2011, 111, 2597–2625.

(2) Scheiner, S. The Hydrogen Bond: A Hundred Years and Counting. Journal of the Indian Institute of Science 2020, 100, 61–76.

(3) Sheu, S. Y.; Yang, D. Y.; Selzle, H. L.; Schlag, E. W. Energetics of hydrogen bonds in peptides. Proceedings of the National Academy of Sciences of the United States of America 2003, 100, 12683–12687.

(4) Herschlag, D.; Pinney, M. M. Hydrogen Bonds: Simple after All? Biochemistry 2018, 57, 3338–3352.

(5) Saunders, L. K.; Nowell, H.; Hatcher, L. E.; Shepherd, H. J.; Teat, S. J.; Allan, D. R.; Raithby, P. R.; Wilson, C. C. Exploring short strong hydrogen bonds engineered in organic acid molecular crystals for temperature dependent proton migration behaviour using single crystal synchrotron X-ray diffraction (SC-SXRD). CrystEngComm 2019, 21, 5249–5260.

(6) Zhou, S.; Wang, L. Unraveling the structural and chemical features of biological short hydrogen bonds. Chemical Science 2019, 10, 7734–7745.

(7) Majerz, I.; Gutmann, M. J. Intermolecular OHN hydrogen bond with a proton moving in 3-methylpyridinium 2,6-dichloro-4-nitrophenolate. RSC Adv. 2015, 5, 95576–95584.

(8) Cleland, W. W.; Kreevoy, M. M. Low-Barrier Hydrogen Bonds and Enzymic Catalysis. Science 1994, 264, 1887–1890.

(9) Perrin, C. L. Are short, low-barrier hydrogen bonds unusually strong? Accounts of Chemical Research 2010, 43, 1550–1557.

(10) Kemp, M. T.; Lewandowski, E. M.; Chen, Y. Low barrier hydrogen bonds in protein structure and function. Biochimica et Biophysica Acta (BBA) - Proteins and Proteomics 2021, 1869, 140557.

(11) Ishikita, H.; Saito, K. Proton transfer reactions and hydrogen-bond networks in protein environments. Journal of The Royal Society Interface 2014, 11, 20130518.

(12) Sigala, P. A.; Ruben, E. A.; Liu, C. W.; Piccoli, P. M. B.; Hohenstein, E. G.; Martínez, T. J.; Schultz, A. J.; Herschlag, D. Determination of Hydrogen Bond Structure in Water versus Aprotic Environments To Test the Relationship Between Length and Stability. Journal of the American Chemical Society 2015, 137, 5730–5740.

(13) Herschlag, D.; Pinney, M. M. Hydrogen Bonds: Simple after All? Biochemistry 2018, 57, 3338–3352.

(14) Chen, D.; Oezguen, N.; Urvil, P.; Ferguson, C.; Dann, S. M.; Savidge, T. C. Regulation of protein-ligand binding affinity by hydrogen bond pairing. Science Advances 2016, 2, e1501240.

(15) Kurczab, R.; Śliwa, P.; Rataj, K.; Kafel, R.; Bojarski, A. J. Salt Bridge in Ligand–Protein Complexes—Systematic Theoretical and Statistical Investigations. Journal of Chemical Information and Modeling 2018, 58, 2224–2238.

(16) Andries, K. et al. A diarylquinoline drug active on the ATP synthase of Mycobacterium tuberculosis. Science 2005, 307, 223–227.

(17) Diacon, A. H. et al. Multidrug-Resistant Tuberculosis and Culture Conversion with Bedaquiline. New England Journal of Medicine 2014, 371, 723–732.

(18) Preiss, L.; Langer, J. D.; Yildiz, Ö.; Eckhardt-Strelau, L.; Guillemont, J. E.; Koul, A.; Meier, T. Structure of the mycobacterial ATP synthase Fo rotor ring in complex with the anti-TB drug bedaquiline. Science Advances 2015, 1, 1–9.

(19) Guo, H.; Courbon, G. M.; Bueler, S. A.; Mai, J.; Liu, J.; Rubinstein, J. L. Structure of mycobacterial ATP synthase bound to the tuberculosis drug bedaquiline. Nature 2021, 589, 143–147.

(20) Haagsma, A. C.; Podasca, I.; Koul, A.; Andries, K.; Guillemont, J.; Lill, H.; Bald, D. Probing the interaction of the diarylquinoline TMC207 with its target mycobacterial ATP synthase. PLoS ONE 2011, 6, 1–7.

(21) Sarathy, J. P.; Ragunathan, P.; Shin, J.; Cooper, C. B.; Upton, A. M.; Grüber, G.; Dick, T. TBAJ-876 Retains Bedaquiline’s Activity against Subunits c and epsilon of Mycobacterium tuberculosis F-ATP Synthase. Antimicrobial Agents and Chemotherapy 2019, 63, 1–11.

(22) E., S.; W., S.; A., N.-C.; V., J.; S., P. New mutations in the mycobacterial ATP synthase: New insights into the binding of the diarylquinoline TMC207 to the ATP synthase C-Ring structure. Antimicrobial Agents and Chemotherapy 2012, 56, 2326–2334.

(23) Meyer, R. L.; Krogsgåard Nielsen, C.; Slavetinsky, C.; Peschel, A.; Ingmer, H.; Vestergaard, M.; Bojer, M. S.; Nøhr-Meldgaard, K. Inhibition of the ATP Synthase Eliminates the Intrinsic Resistance of Staphylococcus aureus towards Polymyxins. mBio 2017, 8, 1–10.

(24) Sarathy, J. P.; Ragunathan, P.; Cooper, C. B.; Upton, A. M.; Grüber, G. TBAJ-876 Displays Bedaquiline-Like Mycobactericidal Potency without Retaining the Parental Drug’s Uncoupler Activity. Antimicrobial Agents and Chemotherapy 2020, 8–13.

(25) Haagsma, A. C.; Abdillahi-Ibrahim, R.; Wagner, M. J.; Krab, K.; Vergauwen, K.; Guillemont, J.; Andries, K.; Lill, H.; Koul, A.; Bald, D. Selectivity of TMC207 towards Mycobacterial ATP synthase compared with that towards the eukaryotic homologue. Antimicrobial Agents and Chemotherapy 2009, 53, 1290–1292.

(26) Balemans, W. et al. Novel antibiotics targeting respiratory ATP synthesis in gram-positive pathogenic bacteria. Antimicrobial Agents and Chemotherapy 2012, 56, 4131–4139.

(27) Melo, M. C.; Bernardi, R. C.; Rudack, T.; Scheurer, M.; Riplinger, C.; Phillips, J. C.; Maia, J. D.; Rocha, G. B.; Ribeiro, J. V.; Stone, J. E.; Neese, F.; Schulten, K.; Luthey-Schulten, Z. NAMD goes quantum: An integrative suite for hybrid simulations. Nature Methods 2018, 15, 351–354.

(28) Mata, I.; Molins, E.; Alkorta, I.; Espinosa, E. Topological Properties of the Electrostatic Potential in Weak and Moderate N…H Hydrogen Bonds. The Journal of Physical Chemistry A 2007, 111, 6425–6433.

(29) Trevisan, L.; Bond, A. D.; Hunter, C. A. Quantitative Measurement of Cooperativity in H-Bonded Networks. Journal of the American Chemical Society 2022, 144, 19499–19507.

(30) Andries, K. et al. A Diarylquinoline Drug Active on the ATP Synthase of !i¿Mycobacterium tuberculosis!/i¿. Science 2005, 307, 223–227.

(31) Petrella, S.; Cambau, E.; Chauffour, A.; Andries, K.; Jarlier, V.; Sougakoff, W. Genetic Basis for Natural and Acquired Resistance to the Diarylquinoline R207910 in Mycobacteria. Antimicrobial Agents and Chemotherapy 2006, 50, 2853–2856.

(32) He, C.; Preiss, L.; Wang, B.; Fu, L.; Wen, H.; Zhang, X.; Cui, H.; Meier, T.; Yin, D. Structural Simplification of Bedaquiline: the Discovery of 3-(4-(N,N-Dimethylaminomethyl)phenyl)quinoline-Derived Antitubercular Lead Compounds. ChemMedChem 2017, 12, 106–119.

(33) Tong, A. S.; Choi, P. J.; Blaser, A.; Sutherland, H. S.; Tsang, S. K.; Guillemont, J.; Motte, M.; Cooper, C. B.; Andries, K.; Van Den Broeck, W.; Franzblau, S. G.; Upton, A. M.; Denny, W. A.; Palmer, B. D.; Conole, D. 6-Cyano Analogues of Bedaquiline as Less Lipophilic and Potentially Safer Diarylquinolines for Tuberculosis. ACS Medicinal Chemistry Letters 2017, 8, 1019–1024.

