## Supplemental Information for "Low-barrier hydrogen bond determines target-binding affinity and specificity of the antitubercular drug bedaquiline"

### SI Methods

#### Simulation systems

The initial structure of the mycobacterial ATP synthase c-ring in complex with bedaquiline (Bq) was taken from the 1.55-Å crystal structure of the complex (PDB id: 4V1F).[1] Using the CHARMM-GUI Membrane Builder,[2–4] the c-ring/Bq complex was embedded in a lipid bilayer oriented perpendicularly with respect to the $z$-axis and composed of 184 palmitoyl-10-methyl-stearoyl-phosphatidylinositol (PTPI), 22 palmitoyl-oleoyl-phosphatidylethanolamine

(POPE), and 34 tetraoleoyl cardiolipin (TOCL) molecules (76.6, 9.2 and 14.2 mol %, respectively), approximating the composition of the inner mycobacterial cell membrane.[5] The system was solvated with TIP3P water molecules in a 103.629 × 103.629 × 101.956 Å rectangular box, and the number of $K^+$ and $Cl^-$ ions was adjusted to maintain a physiological salt concentration of 0.15 M and neutralize the net charge of the system. The CHARMM36m force field was used to describe proteins, lipids, and ions,[6] while the TIP3P model was applied for water. Force fields parameters for bedaquiline (Bq) were obtained from the CHARMM Generalized Force Field (CGenFF),[7] except for four dihedral terms, shown in Fig. S15, which were additionally refined by fitting to the rotational energy profiles computed at the MN12SX/6-31G level using the Gaussian package.[8,9] Bedaquiline partial charges were derived by fitting to electrostatic potential computed at the same level of theory. Two PTPI molecules per leaflet were manually inserted into the c-ring, to seal its central channel, similarly to the previous treatment.[10,11]

The initial structures of the designed Bq analogs bound to the c-ring were prepared by replacing the Bq dimethylamino group in the c-ring/Bq complex by: 2-hydroxy-1-methyl-imidazole (I), formamidic acid (II), and 1-methyl-imidazole (III) substituents.

All systems were subjected to two-step minimization (first keeping all heavy atoms in the c-ring/drug complexes fixed and then without any constraints) followed by initial relaxation using the standard CHARMM-GUI protocol, i.e., a set of short MD runs with progressively weaker restraints on the c-ring/drug atoms. Next, the systems were simulated with the protein backbone atoms harmonically restrained to their initial positions with a force constant of 1000 kJ/(mol·nm$^2$) for 500 ns, and only then equilibrated without any restraints for another 500 ns.

#### Classical molecular dynamics simulations

All classial molecular dynamics (MD) simulations were performed using GROMACS.[12] The simulations were carried out in the isothermal-isobaric (NPT) ensemble. The system temperature was kept at 310 K by Nose-Hoover thermostat.[13] The pressure was maintained semi-isotropically at 1 bar using the Parrinello-Rahman algorithm.[14] Periodic boundary conditions were applied in all three dimension. The electrostatic interactions were evaluated using the Particle Mesh Ewald (PME) algorithm with a real space cut-off radius of 1.2 nm and a grid spacing of 0.12 nm.[15] The dispersion interactions were determined using Lennard-Jones potential with a cut-off radius of 1.2 nm and a force-switch scheme applied over 1.0 to 1.2 nm. The length of all covalent bonds involving a hydrogen atom were constrained using P-LINCS,[16] for the c-ring/drug complexes and lipids, or SETTLE[17] for water. The equations of motion were integrated by the Verlet leapfrog method with a time step of 2 fs.[18] The systems were equilibrated for 500 ns.

To determine the binding free energy Bq to the c-ring using classical MD simulations, we applied the umbrella sampling (US) technique. As a reaction coordinate, we used the center of mass distance between the c-ring backbone heavy atoms and the Bq molecule, $d_{\mathrm{COM}}$ in two different protonation states of E65. To span the range of $d_{\mathrm{COM}}$ corresponding to the full dissociation of Bq from the c-ring (2.5–4.6 nm), we used 21 US windows, separated by 0.1 nm in which the system was restrained by a harmonic potential with a force constant of $500\,\mathrm{kJ/(mol{\cdot}nm^2)}$. The initial configurations for the US windows were taken from an additional non-equilibrium simulation in which Bq was gradually dissociated from the c-ring over 500 ns with a moving harmonic potential with a force constant of $500\,\mathrm{kJ/(mol{\cdot}nm^2)}$. Each of the US windows was simulated for 600 ns and the first 200 ns were discarded as equilibration. Free energy profiles were determined using weighted histogram analysis method (WHAM).[19]

#### Ab initio Molecular Dynamics Simulations

##### Unbiased simulation

QM/MM ab initio molecular dynamics simulations of the above-described systems involving the c-ring/drug complexes embedded in a fully solvated lipid bilayer were performed using

NAMD 2.14[20,21] interfaced with ORCA 4.2.[22] The QM subsystem, depicted in Fig. S4A, included the side chains of two critical acidic residues in the c-ring, E65 and D32, and the dimethylamino group of Bq (32 atoms in total). The alanine side chain was included in the QM subsystem instead of the D32 side chain for the D32A c-ring variant (29 atoms in total; Fig. S4B). The QM subsystem used in the simulations of Bq analogs bound to the wild-type c-ring incorporated the HB-forming functional groups in the ligand molecules and the E65/D32 pair.

The QM forces were calculated using the unrestricted Kohn-Sham scheme with the $\omega$B97X-D3 hybrid functional[23] in combination with the def2-SVP or def2-TZVP basis sets.[24] The simulation was accelerated by applying the resolution-of-identity (RIJ) approximation in combination with the def2/J auxiliary basis set for the evaluation of the Coulomb integrals and the semi-numerical chain-of-spheres algorithm for the Hartree-Fock exchange integrals (COSX).[25] The MM system was described with the CHARMM36m force field[6] for protein, BQ, lipids and ions, and TIP3P model for water. To saturate the covalent bonds between the QM and MM regions, we used hydrogen link atoms with the default charge distribution scheme.[20] Periodic boundary conditions were applied. The electrostatic interaction between the QM and MM systems was modeled through electrostatic embedding, where the MM partial charges surrounding the QM region were passed to ORCA using the default cutoff and charge shifting scheme.[20] Long-range electrostatic interactions between the MM and QM regions, as well as MM electrostatics, were calculated using the Particle Mesh Ewald method (PME) with a real space cutoff of 1.2 nm and a grid spacing of 1.0. The QM partial charges were updated at each step. The Lennard-Jones potential with a cutoff of 1.2 nm was used to describe the intra-MM and QM–MM van der Waals interactions.

The simulations were performed in the NPT ensemble, with the temperature maintained at 310 K using Langevin dynamics with a damping coefficient of 50 ps$^{-1}$. The pressure was maintained at 1 bar using the Langevin piston method with an oscillation period of 0.2 ps and a damping time scale of 0.1 ps. Equations of motion were integrated using the velocity

Verlet algorithm with a time step of 0.5 fs and the total simulation time was 80 ps (for Bq) and 10 ps (for Bq analogs).

##### Free energy simulations

The AIMD-based umbrella sampling technique was applied to determine the changes in the free energy accompanying the proton transfer between the N atom of Bq's amino group and the carboxylate O atom in E65. Separate free energy profiles were obtained for the wild-type c-ring and its D32A variant. The QM/MM setup and MD protocol were the same as those described above for the unbiased simulations. As a reaction coordinate, we used the distance between the carboxylate oxygen atom and the hydrogen atom, $d_{\mathrm{OH}}$. For both WT and D32A, we used 13 equally-spaced US "windows" separated by 0.05 nm, using the harmonic potential with a spring constant of 500 kcal/(mol·nm$^2$) to span the 1.00–1.60 nm range of $d_{\mathrm{OH}}$. The initial configurations for the US windows were taken from additional pulling runs, carried out for WT and D32A separately, in which the proton was displaced from E65 by an externally applied moving harmonic potential with a constant force of 1250 kcal/(mol·nm$^2$). Each of the US windows were simulated for 90 ps and first 40 ps were discarded as equilibration. The free energy profiles were determined using the weighted histogram analysis method (WHAM).[19]

#### Binding energy and hydrogen bond analysis

To calculate the binding energy, $BE$, arising from the hydrogen bonding interaction between the Bq's dimethylamino group and the c-ring's E65 (in the presence or absence of D32), we used the following formula: $BE = E_{\mathrm{complex}} - (E_{\mathrm{Bq}} + E_{\mathrm{protein}})$. Here, $E_{\mathrm{complex}}$ is the energy of the complex involving the entire QM subsystem, $E_{\mathrm{Bq}}$ is the energy of the Bq dimethylamino group, and $E_{\mathrm{protein}}$ is the energy of E65/D32 pair (WT) or E65 alone (D32A mutant). Individual energies were evaluated using density functional theory within Gaussian 16[9] and averaged over 100 random MD frames taken from our unbiased QM/MM AIMD simulations. The QM subsystem taken for these calculations was extracted from the full simulated system and capped with hydrogen atoms. To assess the sensitivity of the binding energy to computational details, the calculations were carried out using three different exchange-correlation functionals, i.e., B3LYP,[26,27] $\omega$B97XD[28] and M06-2X,[29] and two different basis sets, i.e., 6-311++g(d,p) and 6-31g(d). Grimme's D3 dispersion correction was applied for B3LYP and M06-2X and the binding energy was corrected for basis set superposition error (BSSE). For Bq analogs, the identical process was utilized, with the exception that only the B3LYP functional was employed and the energies were averaged over 20 random MD-generated geometries in proportion to the shorter trajectory lengths.

Electron densities at critical points were computed using DAMQT2.0 software[30] and averaged over 100 MD frames randomly extracted from our unbiased QM/MM trajectories.

Table S1: Average binding energy between the Bq's dimethylamino group and E65 in the WT c-ring and its D32A variant (both in the neutral configuration). All values are in kcal/mol.

| XC functional | B3LYP | M06-2X | $\omega$B97XD |
|---|---|---|---|
| 6-311++g(d,p) | | | |
| WT | $-21.48 \pm .77$ | $-20.44 \pm 0.21$ | $-19.17 \pm 0.41$ |
| D32A | $-14.42 \pm .29$ | $-14.64 \pm 0.03$ | $-14.32 \pm 0.15$ |
| 6-31g(d) | | | |
| WT | $-21.60 \pm .43$ | $-23.37 \pm 0.13$ | $-20.19 \pm .41$ |
| D32A | $-14.44 \pm .24$ | $-14.77 \pm 0.03$ | $-16.98 \pm .14$ |

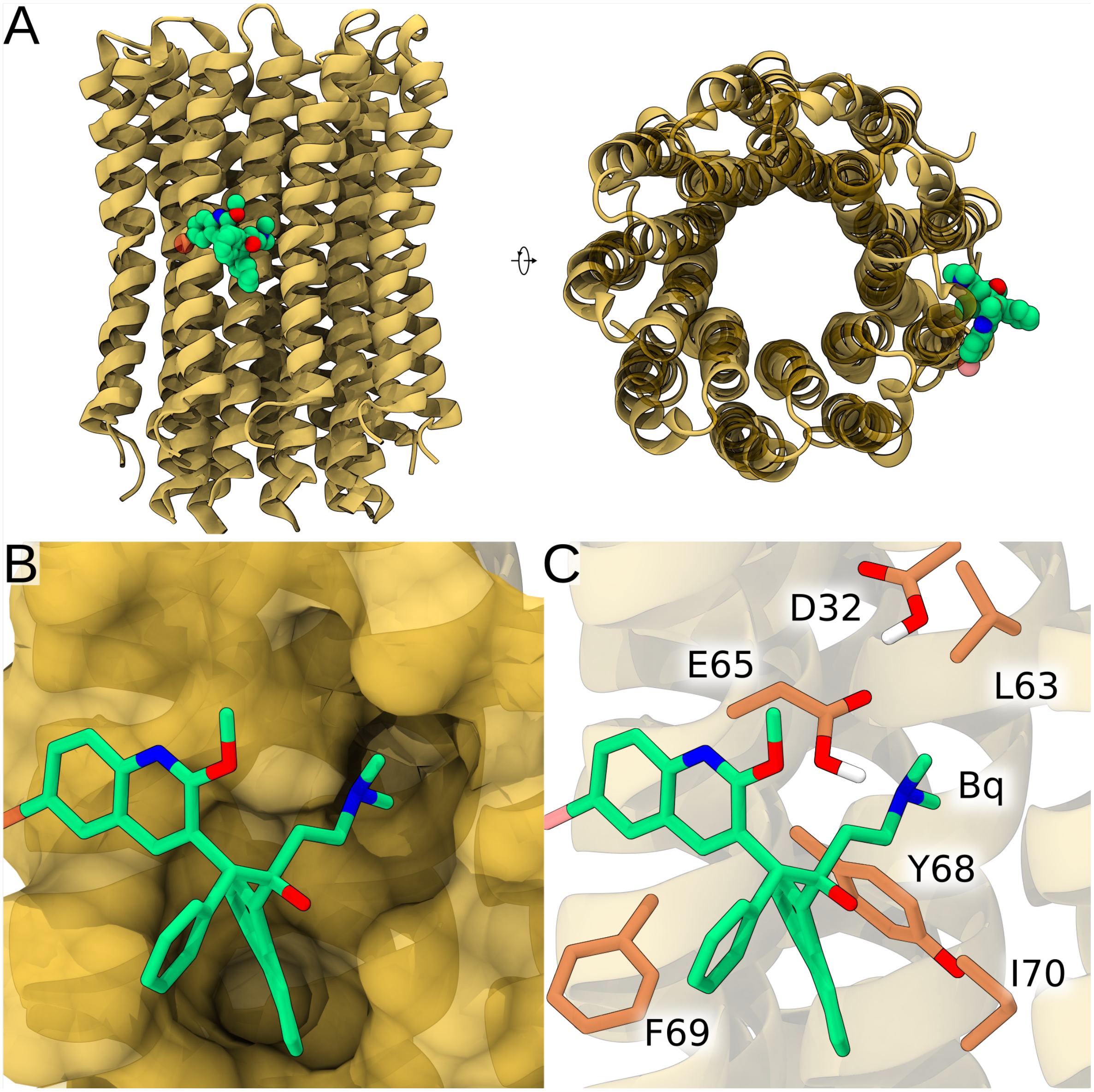


Figure S1: (A) Side and top view of the c-ring (yellow) with bound Bq shown in space-filling representation colored by atom type. (B) Shape complementarity between Bq and its binding pocket in the mycobacterial c-ring. (C) Protein residues forming the Bq-binding pocket in the mycobacterial c-ring.

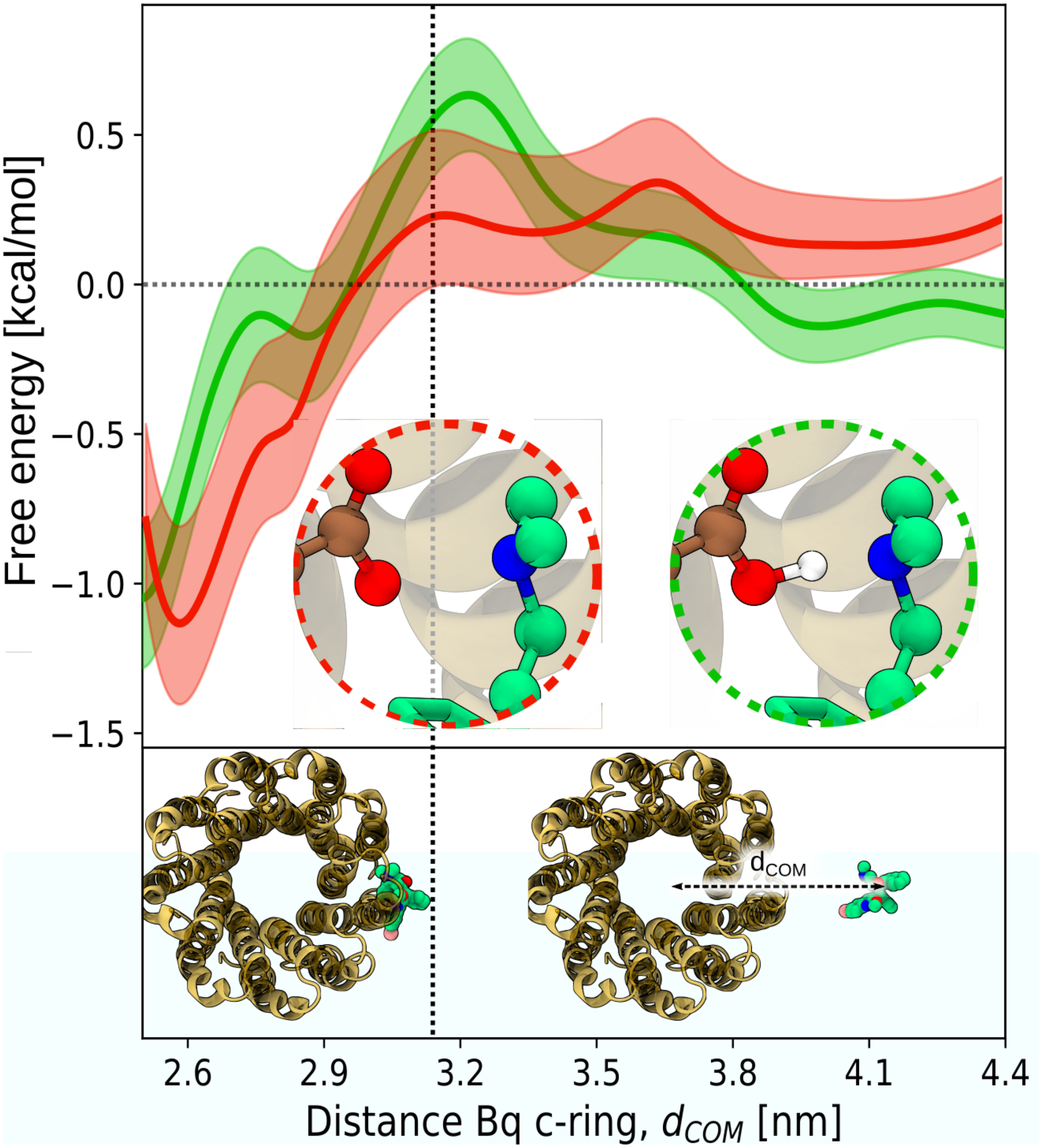


Figure S2: Free energy profile for the Bq binding to the membrane-embedded mycobacterial c-ring, determined with force field-based umbrella sampling simulations. To assess the impact of a direct Bq–E65 hydrogen bond on the binding free energy, the profile was computed twice: once with E65 protonated (green; H-bond present) and once with E65 deprotonated (red; H-bond absent)

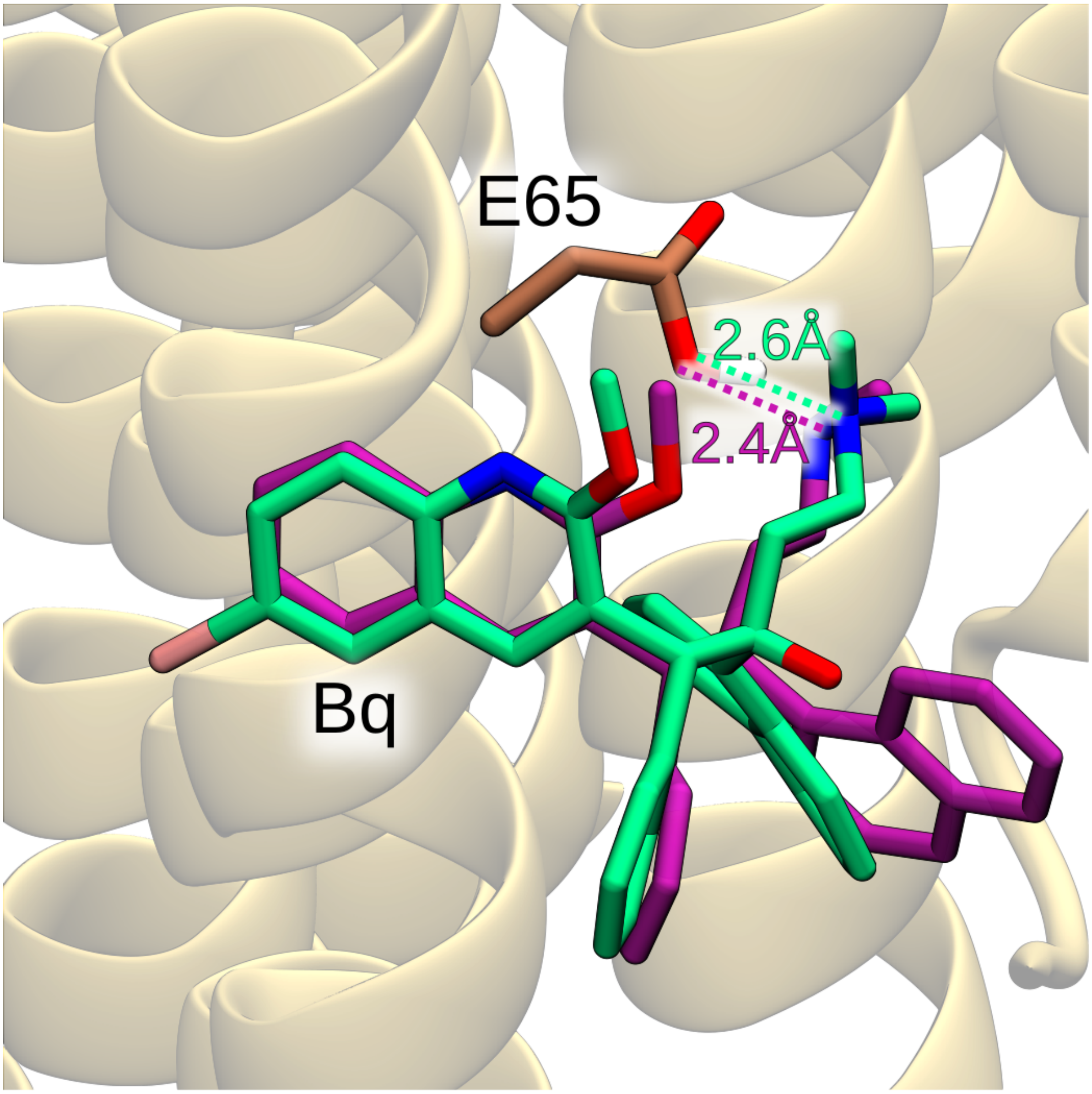


Figure S3: Comparison of the optimal binding pose of Bq identified through force field-based free energy simulations (green) with the crystal structure of the c-ring/Bq complex (PDB id: 4v1f) (purple). The force field's failure to adequately capture the strong nature of hydrogen bonding led to the extension of the Bq–E65 hydrogen bond length to a range commonly observed in hydrogen bonds of moderate strength, measuring 2.6Å.

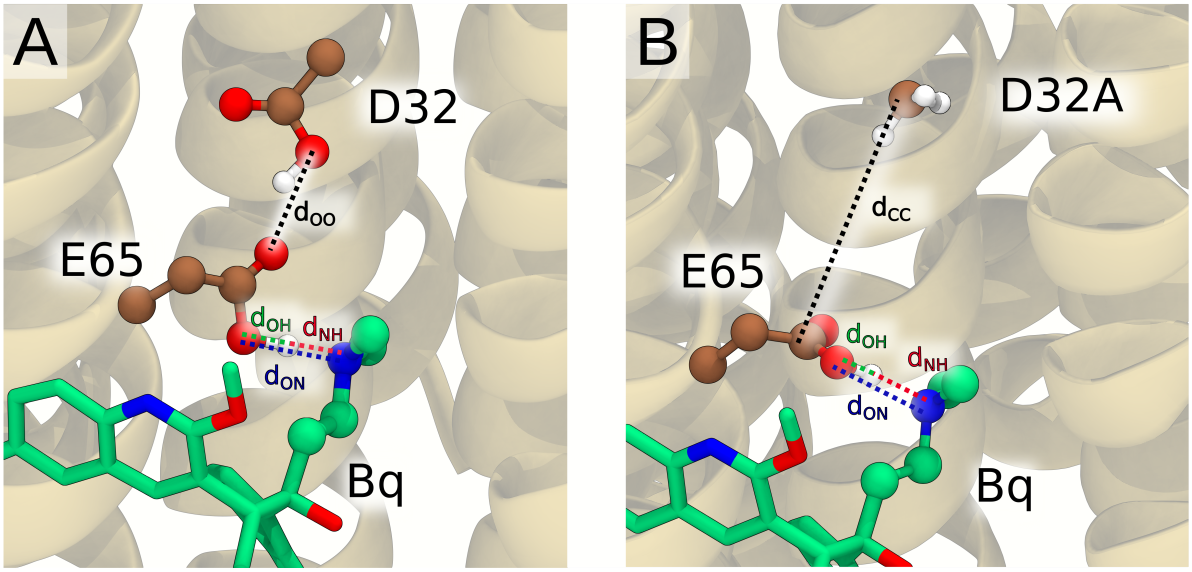


Figure S4: CPK representation depicting the QM subsystem employed in the QM/MM simulations of the wild-type c-ring (A) or its D32A variant (B) in complex with Bq. Non-polar hydrogen atoms were omitted for the sake of clarity. Additionally, the distances considered in this work are defined.

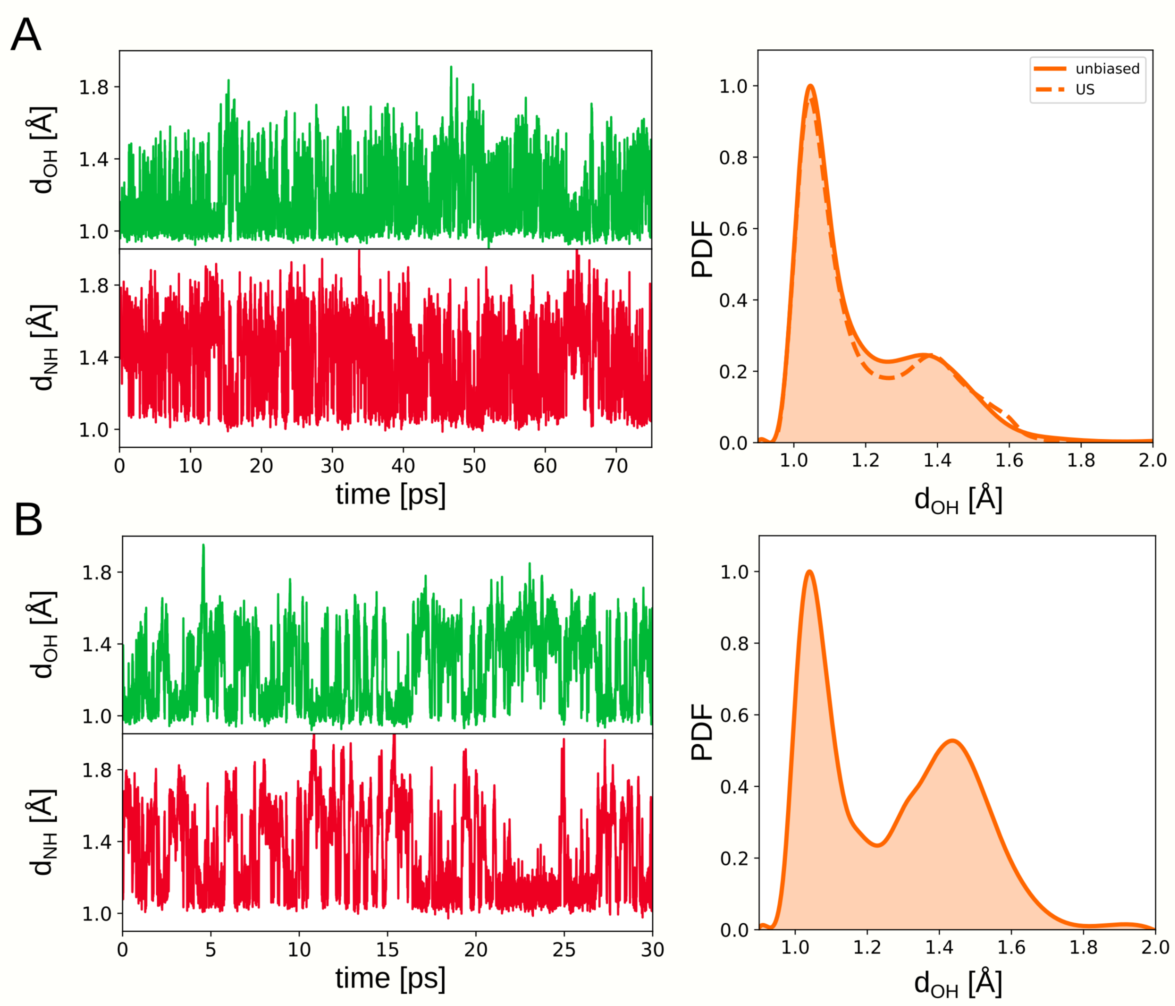


Figure S5: Effect of the basis set employed on the dynamic behavior of a proton involved in the hydrogen bond formed between Bq and E65 the the wild-type c-ring. (left) Time evolution of the $d_{\mathrm{OH}}$ and $d_{\mathrm{NH}}$ distances in the unbiased simulation carried out using double-zeta basis set (A) and triple-zeta basis set (B). (right) Comparison of the probability density of finding the proton at different distances from the carboxylate O atom of E65 derived from the unbiased (solid line) and umbrella simulations (dashed line) using a double-zeta (A) and triple-zeta (B) basis set.

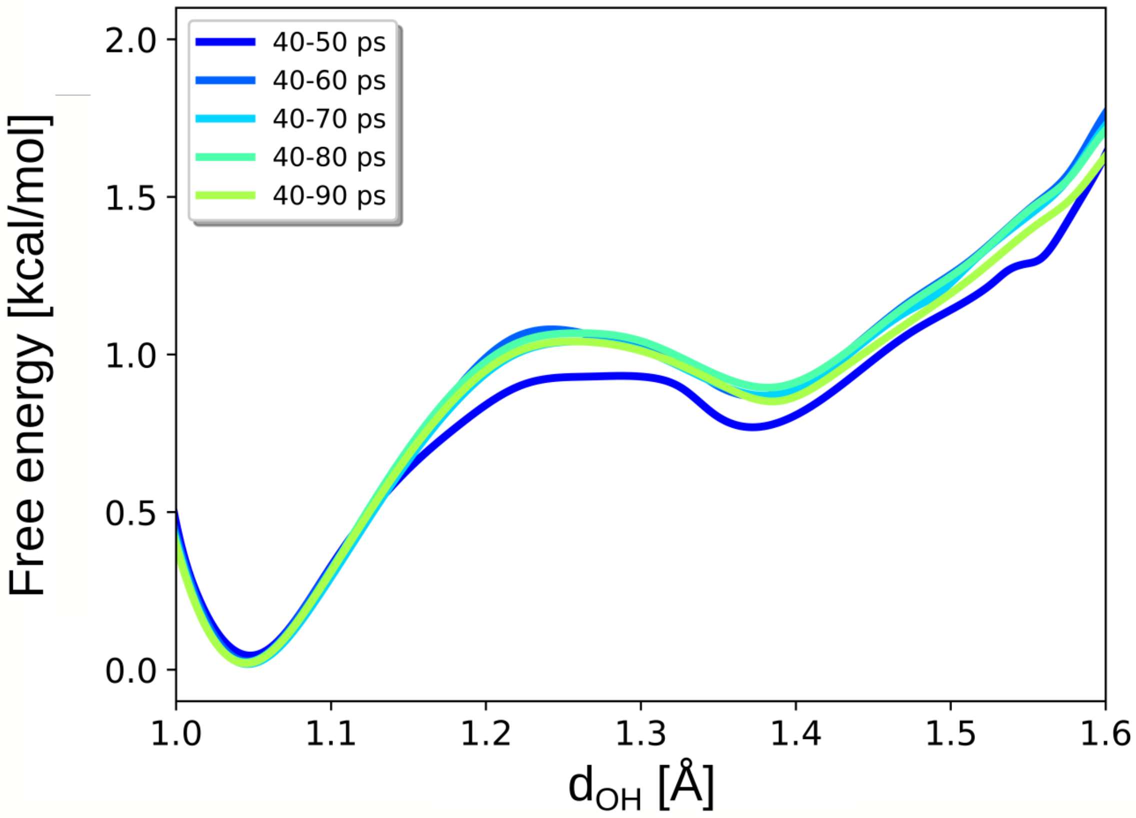


Figure S6: Convergence of the free energy profiles for the proton transfer between heteroatoms involved in the hydrogen bond between Bq and E65 in the WT c-ring as a function of the length of US trajectories used for analysis.

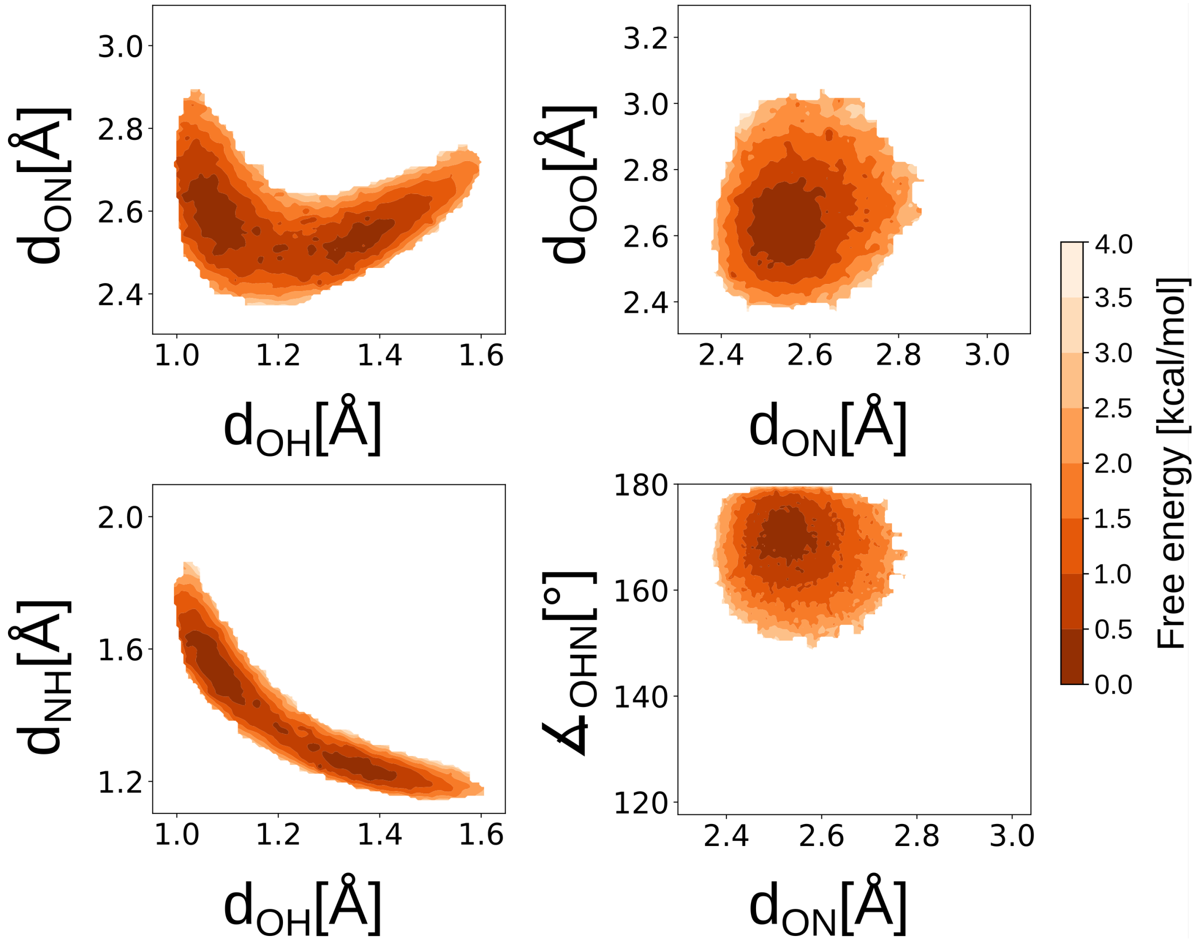


Figure S7: Free energy landscapes describing the correlations between the coordinates defining the geometry of the H-bond between Bq and E65 in the wild type c-ring: the distances $d_{\mathrm{OH}}$, $d_{\mathrm{ON}}$, $d_{\mathrm{NH}}$, $d_{\mathrm{OO}}$ (see: Fig. S4), and the angle $\angle_{\mathrm{OHN}}$).

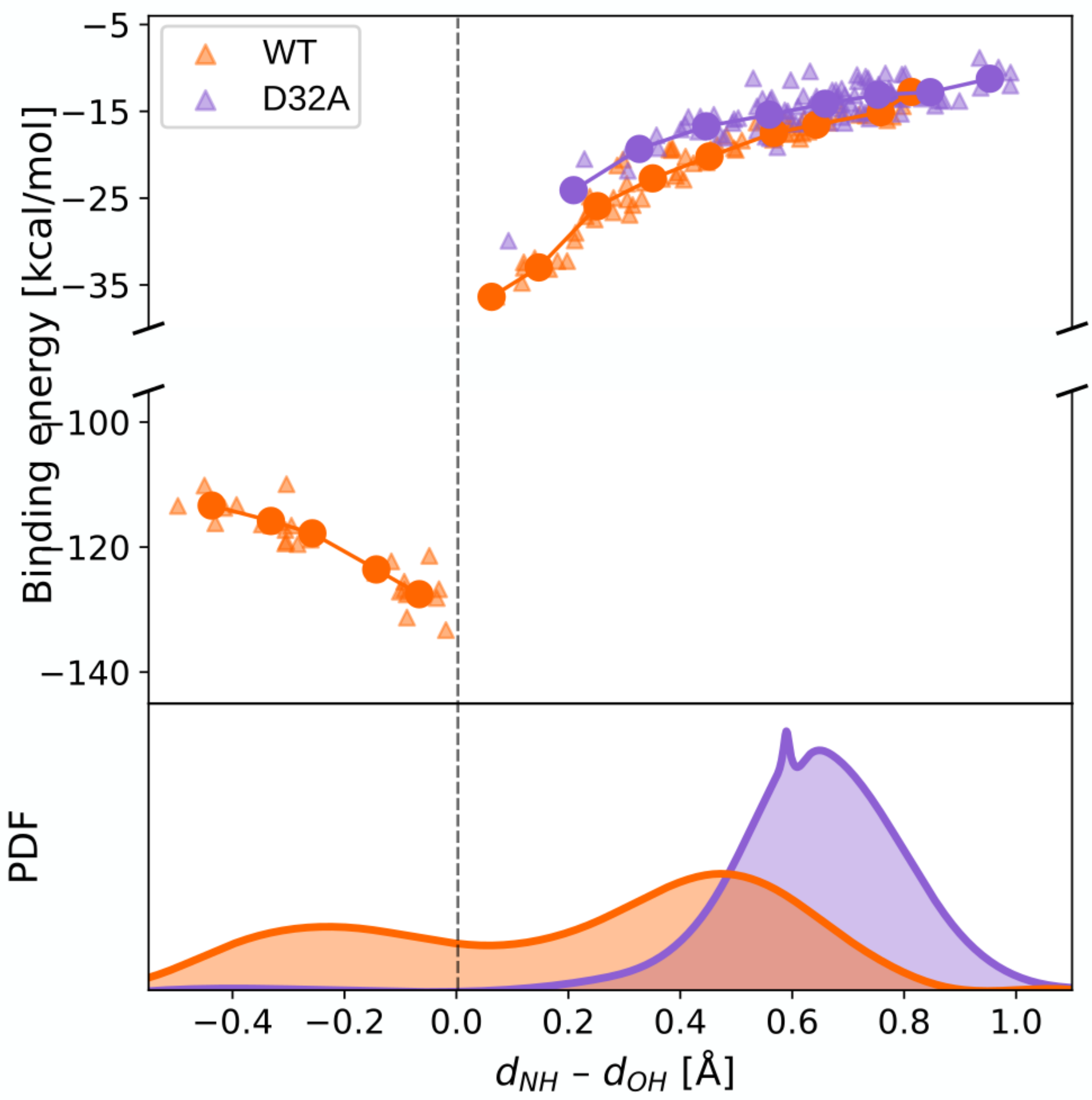


Figure S8: Binding energy arising from the hydrogen bonding interaction between the Bq's dimethylamino group and the c-ring's E65, in the presence (orange) and absence (purple) of D23, as a function of the proton transfer between the two heteroatoms. Since the proton transfer coordinate is defined as the distance difference $d_{\mathrm{NH}} - d_{\mathrm{OH}}$, its negative values correspond to the proton located on Bq (zwitterionic or 'salt-bridge' configuration) and positive values correspond to the proton located on the c-ring (neutral configuration). Triangles represent individual binding energies computed for 100 geometries extracted randomly from the unbiased QM/MM simulations (see SI Methods). The solid line shows the running average, calculated separately for the neutral and zwitterionic configurations. The electrostatic attraction between oppositely-charged ions is responsible for a significant portion of the binding energy obtained in the zwitterionic configuration. In fact, when two elementary charges are separated by a distance of 3.2Å (representing the average distance between the centers of charge of the Bq's dimethylamino group and E65 carboxylic group), the electrostatic interaction energy is approximately 100 kcal/mol. The distributions presented at the bottom depict the probabilities of finding the system at different values of $d_{\mathrm{NH}} - d_{\mathrm{OH}}$, as derived from our unbiased QM/MM simulations.

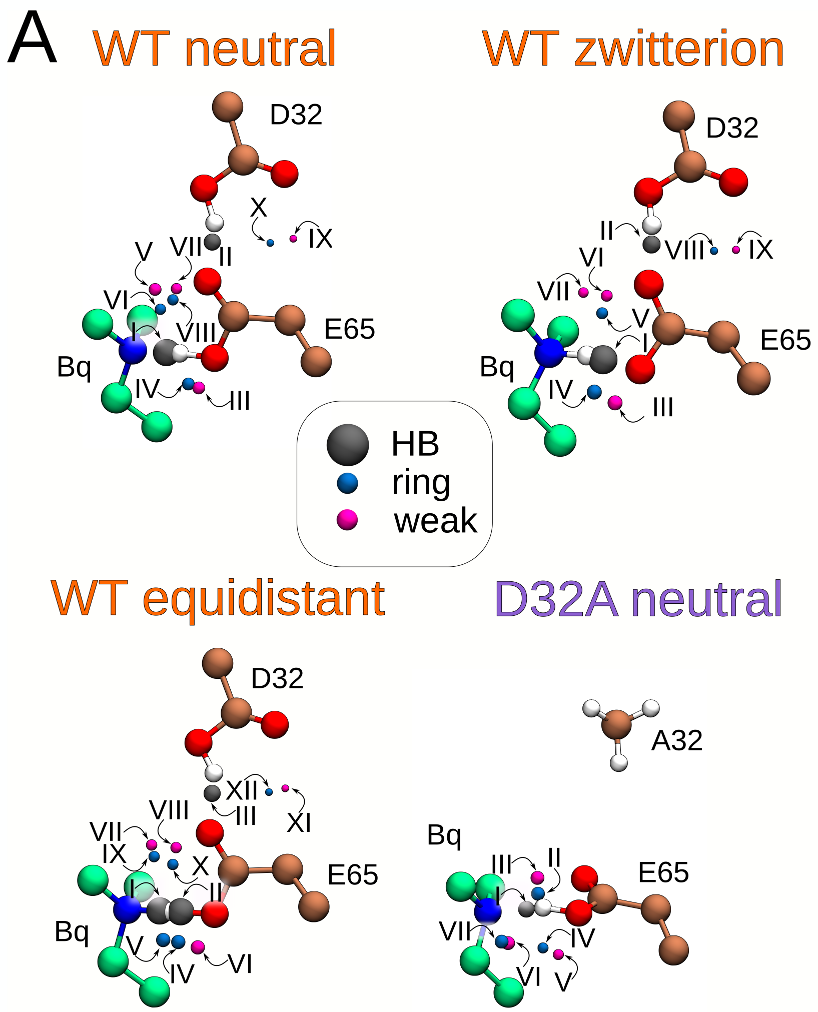


B

**WT neutral**

| CP | ρ |
|---|---|
| I | 0.0967 |
| II | 0.0491 |
| III | 0.0096 |
| IV | 0.0099 |
| V | 0.0091 |
| VI | 0.0067 |
| VII | 0.0052 |
| VIII | 0.0054 |
| IX | 0.0018 |
| X | 0.0023 |

**WT zwitterion**

| CP | ρ |
|---|---|
| I | 0.1020 |
| II | 0.0450 |
| III | 0.0113 |
| IV | 0.0152 |
| V | 0.0064 |
| VI | 0.0077 |
| VII | 0.0047 |
| VIII | 0.0020 |
| IX | 0.0025 |

**WT equidistant**

| CP | ρ |
|---|---|
| I | 0.1641 |
| II | 0.1581 |
| III | 0.0403 |
| IV | 0.0119 |
| V | 0.0124 |
| VI | 0.0138 |
| VII | 0.0052 |
| VIII | 0.0070 |
| IX | 0.0050 |
| X | 0.0059 |
| XI | 0.1588 |
| XII | 0.1588 |

**D32A neutral**

| CP | ρ |
|---|---|
| I | 0.0627 |
| II | 0.0104 |
| III | 0.0167 |
| IV | 0.0065 |
| V | 0.0072 |
| VI | 0.0111 |
| VII | 0.0127 |

Figure S9: (A) Positions of critical points corresponding to different types of interactions at the interface between the Bq amino group and the c-ring (legend: HB – hydrogen-bond critcal point, ring – ring critical point, weak – weak-interaction critical point). Four different cases were considered: WT c-ring with a proton located on E65 (WT neutral), WT c-ring with a proton located on Bq (WT zwitterion), WT c-ring with a proton equidistant from both heteroatoms (WT equidistant) and D32A variant of the c-ring with a proton located on E65 (D32A neutral). The sizes of the spheres are proportional to the corresponding values of electron density, which are explicitly shown in panel (B) in e/Å$^3$. We averaged both the positions of critical points and their electron densities across a set of 100 random MD frames extracted from our unbiased QM/MM trajectories.

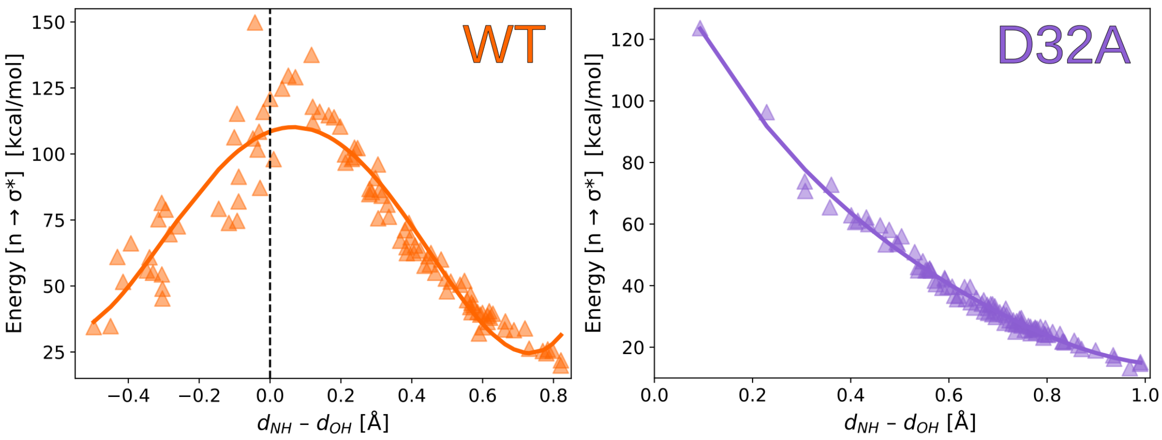


Figure S10: Computed Natural Bond Orbital (NBO) $2^{\mathrm{nd}}$-order estimate of $\mathrm{n}_D \rightarrow \sigma^*_{AH}$ interaction energy for the Bq–E65 H-bond as function of the distance difference $d_{\mathrm{NH}} - d_{\mathrm{OH}}$ (see caption for Fig. S8). The NBO analysis was performed at the B3LYP/6-311++g(d,p) level of theory.

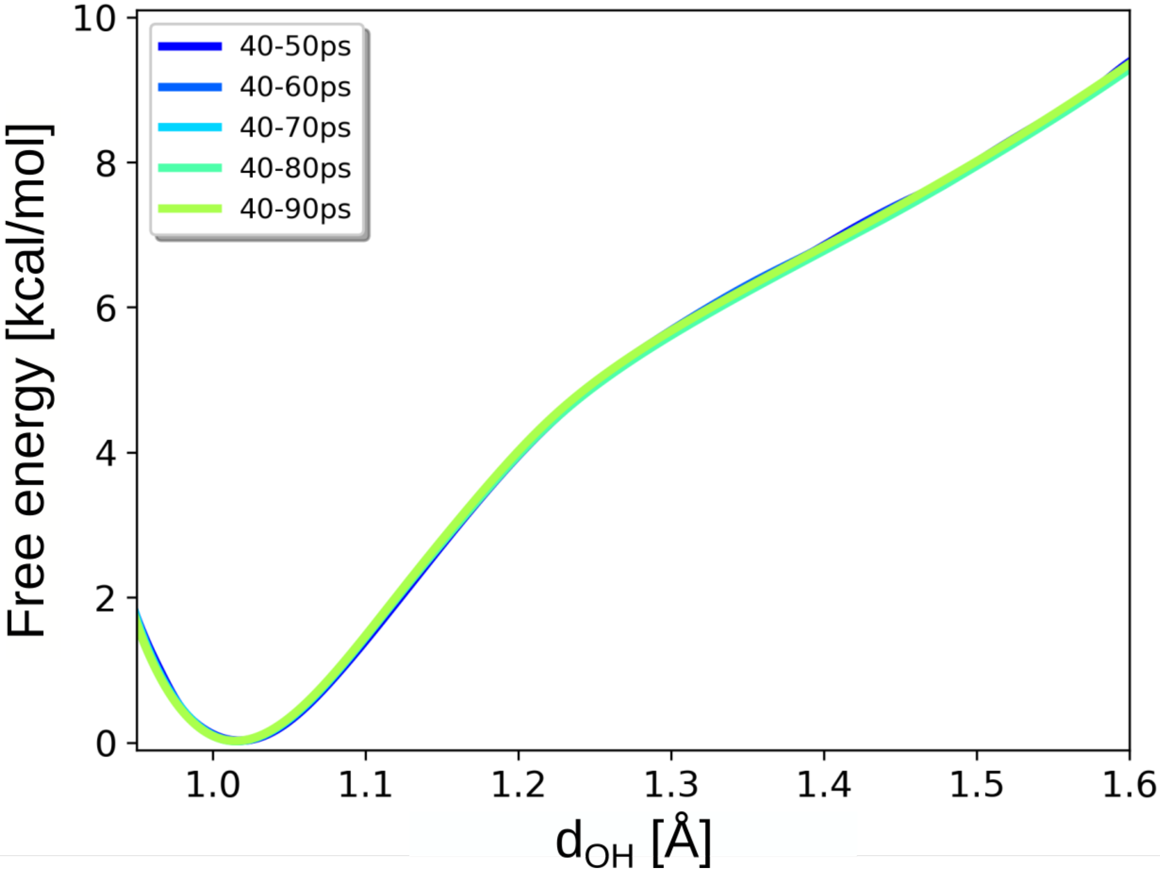


Figure S11: Convergence of the free energy profiles for the proton transfer between heteroatoms involved in the hydrogen bond between Bq and E65 in D32A variant of the c-ring as a function of the length of US trajectories used for analysis.

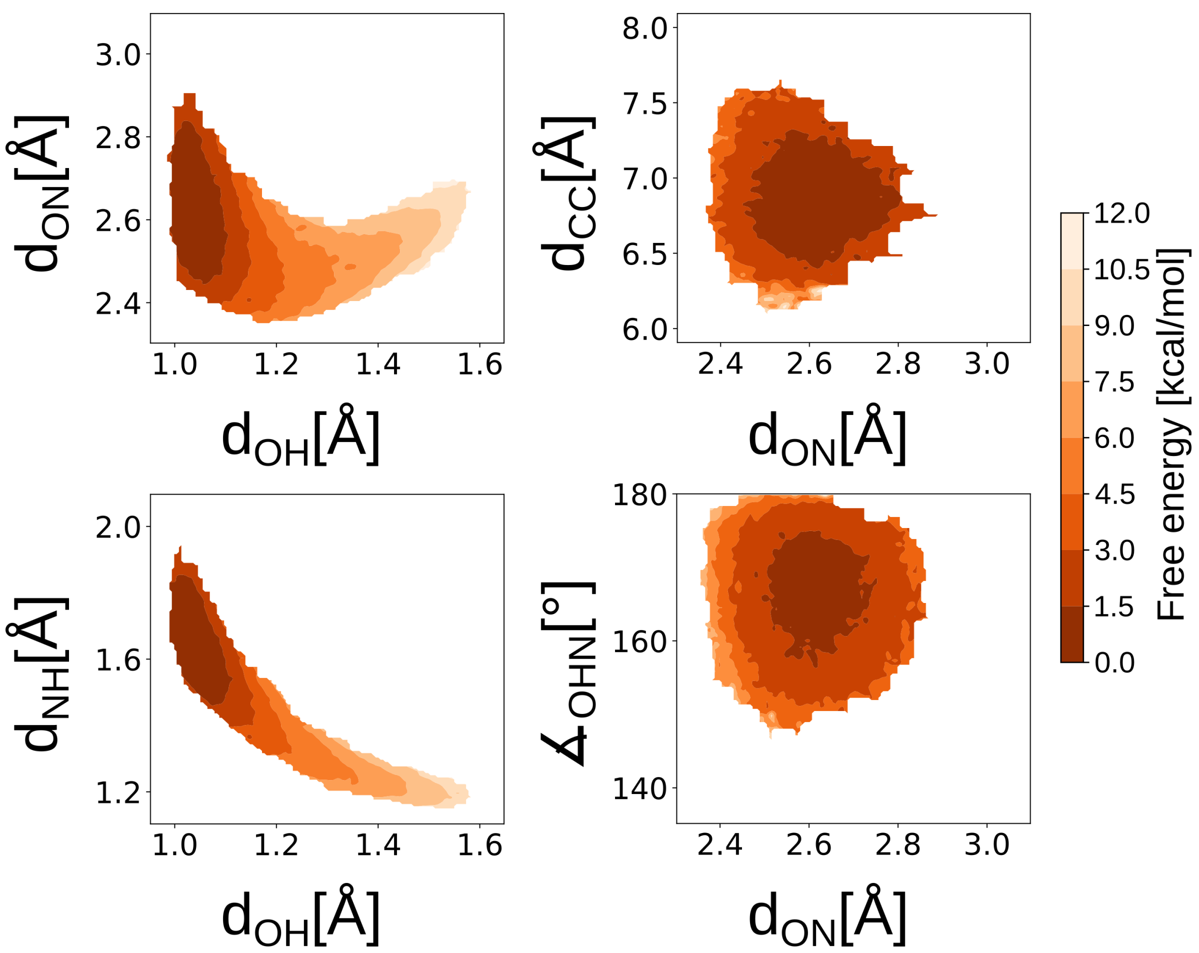


Figure S12: Free energy landscapes describing the correlations between the coordinates defining the geometry of the H-bond between Bq and E65 in D32A variant of the c-ring.: the distances $d_{OH}$, $d_{ON}$, $d_{NH}$, $d_{OO}$ (see: Fig. S4), and the angle $\angle_{OHN}$).

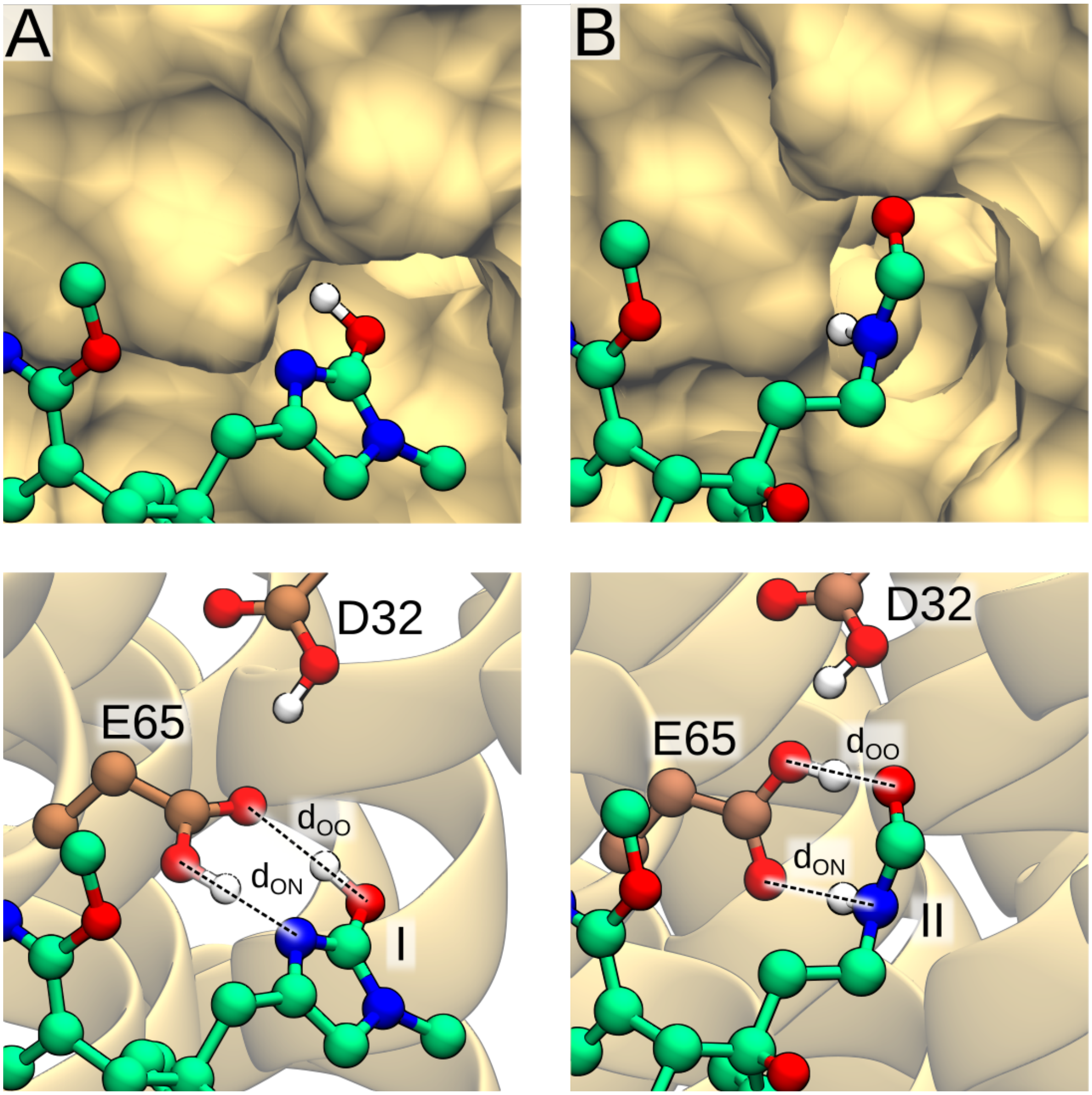


Figure S13: Hydrogen bonding interaction between Bq analogs (I and II) and the wild-type c-ring as identified by our QM/MM simulations. The average heavy-atom distances for the ligand–c-ring hydrogen bonds are: I – $d_{ON} = 2.48$ Å, $d_{OO} = 2.76$ Å, II – $d_{ON} = 2.70$ Å, $d_{OO} = 2.45$ Å,

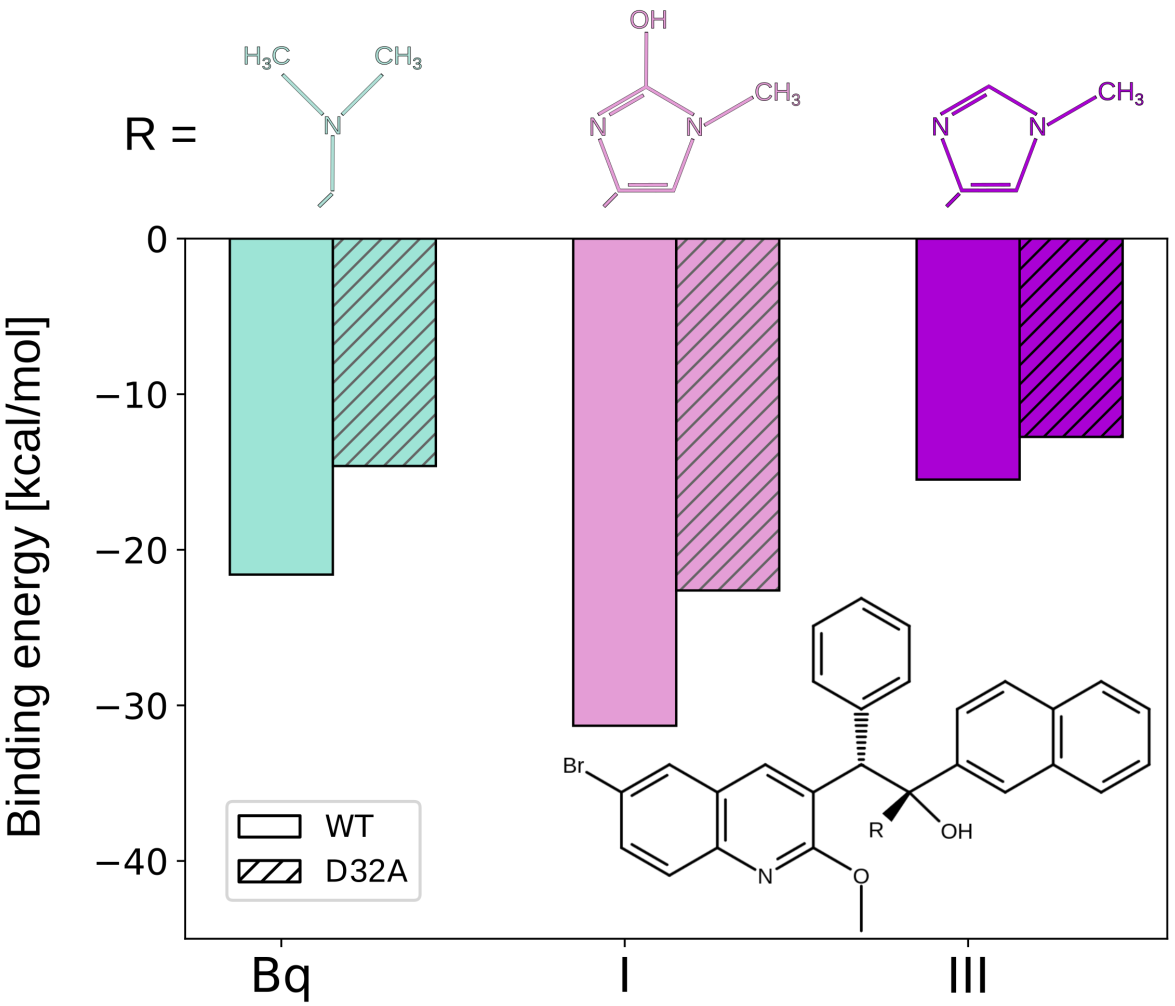


Figure S14: Hydrogen bond strength between Bq or its two analogs (I/III) and E65 in the WT c-ring and its D32A variant, quantified as the binding energy between the ligand's HB-forming group (R) and either E65/D32 pair (WT) or E65 alone (D32A).

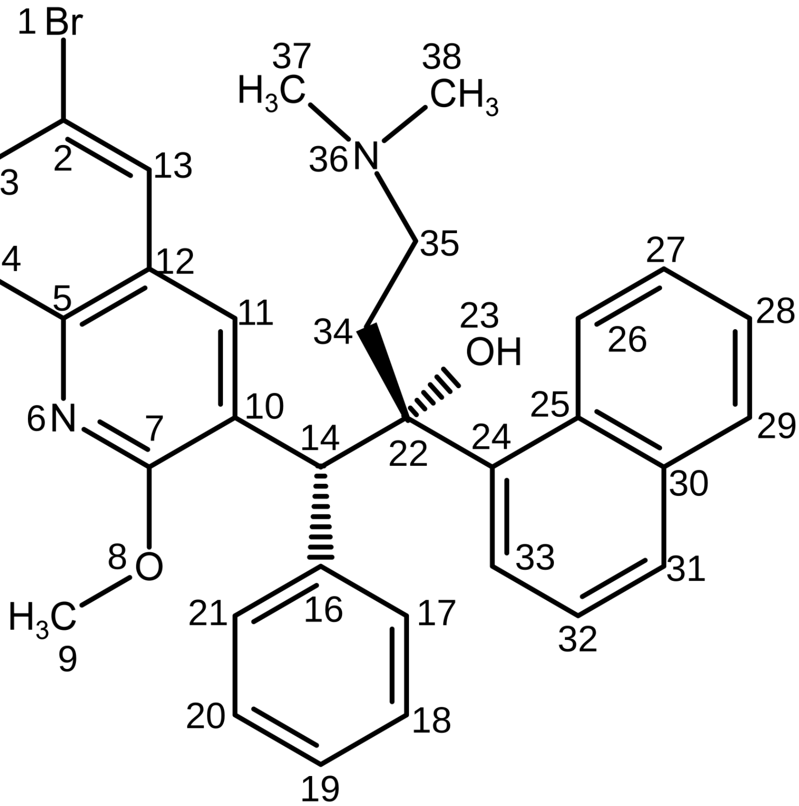


| 8 | 7 | 6 | 5 | phi0 | kphi | multiplicity |
|---|---|---|---|---|---|---|
| OG301 | CG2R61 | NG2R60 | CG2R61 | 180.00 | 34.158176 | 1 |
| OG301 | CG2R61 | NG2R60 | CG2R61 | 0.00 | 13.710968 | 2 |
| OG301 | CG2R61 | NG2R60 | CG2R61 | 0.00 | 18.463992 | 3 |
| OG301 | CG2R61 | NG2R60 | CG2R61 | 0.00 | 18.38868 | 4 |
| OG301 | CG2R61 | NG2R60 | CG2R61 | 180.00 | 22.873928 | 6 |
| **6** | **7** | **8** | **9** | | | |
| NG2R60 | CG2R61 | OG301 | CG331 | 180.00 | 0.217568 | 1 |
| NG2R60 | CG2R61 | OG301 | CG331 | 180.00 | 19.459784 | 2 |
| NG2R60 | CG2R61 | OG301 | CG331 | 0.00 | 2.46856 | 3 |
| NG2R60 | CG2R61 | OG301 | CG331 | 180.00 | 2.11292 | 4 |
| NG2R60 | CG2R61 | OG301 | CG331 | 0.00 | 1.577368 | 6 |
| **24** | **22** | **14** | **10** | | | |
| CG2R61 | CG301 | CG311 | CG2R61 | 180.00 | 4.861808 | 2 |
| CG2R61 | CG301 | CG311 | CG2R61 | 180.00 | 3.602424 | 3 |
| CG2R61 | CG301 | CG311 | CG2R61 | 0.00 | 3.74468 | 4 |
| CG2R61 | CG301 | CG311 | CG2R61 | 180.00 | 1.502056 | 6 |
| **34** | **35** | **36** | **37** | | | |
| CG321 | CG321 | NG301 | CG331 | 0.00 | 3.7656 | 1 |
| CG321 | CG321 | NG301 | CG331 | 0.00 | 0.799144 | 2 |
| CG321 | CG321 | NG301 | CG331 | 0.00 | 5.39736 | 3 |
| CG321 | CG321 | NG301 | CG331 | 0.00 | 0.677808 | 4 |
| CG321 | CG321 | NG301 | CG331 | 0.00 | 0.255224 | 6 |

Figure S15: Force field parameters for the four refined dihedral terms.

### References


(1) Preiss, L.; Langer, J. D.; Yildiz, Ö.; Eckhardt-Strelau, L.; Guillemont, J. E.; Koul, A.; Meier, T. Structure of the mycobacterial ATP synthase Fo rotor ring in complex with the anti-TB drug bedaquiline. *Science Advances* **2015**, *1*, 1–9.

(2) Jo, S.; Kim, T.; Iyer, V. G.; Im, W. CHARMM-GUI: A web-based graphical user interface for CHARMM. *Journal of Computational Chemistry* **2008**, *29*, 1859–1865.

(3) Wu, E. L.; Cheng, X.; Jo, S.; Rui, H.; Song, K. C.; Dávila-Contreras, E. M.; Qi, Y.; Lee, J.; Monje-Galvan, V.; Venable, R. M.; Klauda, J. B.; Im, W. CHARMM-GUI Membrane Builder toward realistic biological membrane simulations. *Journal of Computational Chemistry* **2014**, *35*, 1997–2004.

(4) Lee, J. et al. CHARMM-GUI Input Generator for NAMD, GROMACS, AMBER, OpenMM, and CHARMM/OpenMM Simulations Using the CHARMM36 Additive Force Field. *Journal of Chemical Theory and Computation* **2016**, *12*, 405–413.

(5) Bansal-Mutalik, R.; Nikaido, H. Mycobacterial outer membrane is a lipid bilayer and the inner membrane is unusually rich in diacyl phosphatidylinositol dimannosides. *Proceedings of the National Academy of Sciences* **2014**, *111*, 4958–4963.

(6) Huang, J.; Rauscher, S.; Nawrocki, G.; Ran, T.; Feig, M.; de Groot, B. L.; Grubmüller, H.; MacKerell, A. D. CHARMM36m: an improved force field for folded and intrinsically disordered proteins. *Nature Methods* **2017**, *14*, 71–73.

(7) Vanommeslaeghe, K.; Hatcher, E.; Acharya, C.; Kundu, S.; Zhong, S.; Shim, J.; Darian, E.; Guvench, O.; Lopes, P.; Vorobyov, I.; Mackerell Jr., A. D. CHARMM general force field: A force field for drug-like molecules compatible with the CHARMM all-atom additive biological force fields. *Journal of Computational Chemistry* **2010**, *31*, 671–690.

(8) Peverati, R.; Truhlar, D. G. Screened-exchange density functionals with broad accuracy for chemistry and solid-state physics. *Phys. Chem. Chem. Phys.* **2012**, *14*, 16187–16191.

(9) Frisch, M. e.; Trucks, G.; Schlegel, H.; Scuseria, G.; Robb, M.; Cheeseman, J.; Scalmani, G.; Barone, V.; Petersson, G.; Nakatsuji, H., et al. Gaussian 16. 2016.

(10) Zhou, W.; Marinelli, F.; Nief, C.; Faraldo-Gómez, J. D. Atomistic simulations indicate the c-subunit ring of the F1Fo ATP synthase is not the mitochondrial permeability transition pore. *eLife* **2017**, *6*.

(11) Marciniak, A.; Chodnicki, P.; Hossain, K. A.; Slabonska, J.; Czub, J. Determinants of Directionality and Efficiency of the ATP Synthase Fo Motor at Atomic Resolution. *The Journal of Physical Chemistry Letters* **2022**, *13*, 387–392.

(12) Van Der Spoel, D.; Lindahl, E.; Hess, B.; Groenhof, G.; Mark, A. E.; Berendsen, H. J. C. GROMACS: Fast, flexible, and free. *Journal of Computational Chemistry* **2005**, *26*, 1701–1718.

(13) Nosé, S. A unified formulation of the constant temperature molecular dynamics methods. *The Journal of Chemical Physics* **1984**, *81*, 511–519.

(14) Parrinello, M.; Rahman, A. Polymorphic transitions in single crystals: A new molecular dynamics method. *Journal of Applied Physics* **1981**, *52*, 7182–7190.

(15) Essmann, U.; Perera, L.; Berkowitz, M. L.; Darden, T.; Lee, H.; Pedersen, L. G. A smooth particle mesh Ewald method. *The Journal of Chemical Physics* **1995**, *103*, 8577–8593.

(16) Hess, B. P-LINCS: A Parallel Linear Constraint Solver for Molecular Simulation. *Journal of Chemical Theory and Computation* **2008**, *4*, 116–122.

(17) Miyamoto, S.; Kollman, P. A. Settle: An analytical version of the SHAKE and RATTLE

algorithm for rigid water models. *Journal of Computational Chemistry* **1992**, *13*, 952–962.

(18) Verlet, L. Computer "Experiments" on Classical Fluids. I. Thermodynamical Properties of Lennard-Jones Molecules. *Phys. Rev.* **1967**, *159*, 98–103.

(19) Kumar, S.; Rosenberg, J. M.; Bouzida, D.; Swendsen, R. H.; Kollman, P. A. Multidimensional free-energy calculations using the weighted histogram analysis method. *Journal of Computational Chemistry* **1995**, *16*, 1339–1350.

(20) Melo, M. C.; Bernardi, R. C.; Rudack, T.; Scheurer, M.; Riplinger, C.; Phillips, J. C.; Maia, J. D.; Rocha, G. B.; Ribeiro, J. V.; Stone, J. E.; Neese, F.; Schulten, K.; Luthey-Schulten, Z. NAMD goes quantum: An integrative suite for hybrid simulations. *Nature Methods* **2018**, *15*, 351–354.

(21) Phillips, J. C. et al. Scalable molecular dynamics on CPU and GPU architectures with NAMD. *The Journal of Chemical Physics* **2020**, *153*, 044130.

(22) Neese, F. The ORCA program system. *WIREs Computational Molecular Science* **2012**, *2*, 73–78.

(23) Lin, Y.-S.; Li, G.-D.; Mao, S.-P.; Chai, J.-D. Long-Range Corrected Hybrid Density Functionals with Improved Dispersion Corrections. *Journal of Chemical Theory and Computation* **2013**, *9*, 263–272.

(24) Grimme, S. Density functional theory with London dispersion corrections. *Wiley Interdisciplinary Reviews: Computational Molecular Science* **2011**, *1*, 211–228.

(25) Kossmann, S.; Neese, F. Comparison of two efficient approximate Hartee–Fock approaches. *Chemical Physics Letters* **2009**, *481*, 240–243.

(26) Becke, A. D. Density-functional thermochemistry. III. The role of exact exchange. *The Journal of Chemical Physics* **1993**, *98*, 5648–5652.

(27) Lee, C.; Yang, W.; Parr, R. G. Development of the Colle-Salvetti correlation-energy formula into a functional of the electron density. *Phys Rev B Condens Matter* **1988**, *37*, 785–789.

(28) Chai, J.-D.; Head-Gordon, M. Long-range corrected hybrid density functionals with damped atom–atom dispersion corrections. *Phys. Chem. Chem. Phys.* **2008**, *10*, 6615–6620.

(29) Zhao, Y.; Truhlar, D. G. The M06 suite of density functionals for main group thermochemistry, thermochemical kinetics, noncovalent interactions, excited states, and transition elements: two new functionals and systematic testing of four M06-class functionals and 12 other functionals. *Theoretical Chemistry Accounts* **2008**, *120*, 215–241.

(30) Kumar, A.; Yeole, S. D.; Gadre, S. R.; López, R.; Rico, J. F.; Ramírez, G.; Ema, I.; Zorrilla, D. DAMQT 2.1. 0: a new version of the DAMQT package enabled with the topographical analysis of electron density and electrostatic potential in molecules. 2015.
